# A computational Evo-Devo approach for elucidating the roles of PLETHORA transcription factors in regulating root development

**DOI:** 10.1101/2024.12.03.626639

**Authors:** Joel Rodríguez Herrera, Kenia Aislinn Galván Alcaraz, Ramsés Uriel Albarrán Hernández, Juan Pablo Villa Núñez, Gustavo Rodríguez Alonso, Svetlana Shishkova

**Affiliations:** Departamento de Biología Molecular de Plantas, Instituto de Biotecnología, Universidad Nacional Autónoma de México. Cuernavaca, Morelos, México. C.P. 62210; Centro de Investigación en Dinámica Celular, Instituto de Investigación en Ciencias Básicas y Aplicadas. Universidad Autónoma del Estado de Morelos. Cuernavaca, Morelos, México. C.P. 62210

## Abstract

PLETHORA (PLT) transcription factors play essential roles in regulating various developmental processes in plants, including embryogenesis, rhizotaxis, phyllotaxis, and most prominently, root development, by regulating cell proliferation and differentiation along the root. Despite their important roles in plant development, PLT transcription factors have mainly been studied in *Arabidopsis thaliana* and only a few crop species. *A. thaliana* has six *PLT* genes, which are expressed in overlapping domains and have partially redundant activities, with numerous shared target genes. Here we identified putative *PLT* orthologs across Viridiplantae, including representatives of all extant plant clades, reconstructing the molecular phylogeny of PLTs and integrating synteny and microsynteny analyses. We suggest that PLTs arose by neofunctionalization prior to the divergence of Spermatophyta and that they might regulate their target genes in a context-specific manner given the presence of intrinsically disordered regions at their N- and C-termini. After identifying direct PLT targets in public databases, we inferred a gene regulatory network driven by PLTs in the root apical meristem in six angiosperm species. Our results suggest that the direct PLT targets regulate ribosome and ribonucleoprotein biogenesis as well as RNA processing, among other basic cellular processes. The central relevance of these processes may account for the high conservation and stability of PLT-driven gene regulatory networks across angiosperms.

## Introduction

The evolution and adaptation of plants have been greatly influenced by the flexibility of their molecular responses to developmental and environmental cues. Furthermore, the morphological changes and radiation patterns driven by natural selection usually coincide with changes in the transcriptional landscapes of organisms (1). Transcription factors (TFs) are key to the remodeling of transcriptional landscapes; these DNA binding proteins recognize short, specific *cis*-elements to regulate the transcription of their target genes. TFs play important roles in orchestrating developmental and biotic/abiotic stress responses in plants, thereby underlying plant adaptability and evolution (2). Plant TF families have undergone numerous origination, expansion, and contraction events over the course of their evolution (3). Our understanding of plant TF evolution has deeply benefited from genome sequencing projects and transcriptome analysis (3,4).

One of the largest plant TF families, the APETALA 2/ETHYLENE-RESPONSIVE ELEMENT BINDING PROTEIN (AP2/EREBP) family, is characterized by the presence of an AP2 DNA-binding domain (5,6). The AP2/EREBP family is divided into three subfamilies, the EREBP, RAV (RELATED TO ABI3/VP1), and AP2 subfamilies, based on the number of AP2 domains and whether they include additional domains (7,8). AP2 subfamily proteins contain two AP2 domains, which are ∼60 amino acids long and separated by a linker of ∼25 amino acids. The AP2 subfamily is further divided into the euAP2 and AINTEGUMENTA (ANT) clades, and the ANT clade is divided into three major groups: the basal ANT, euANT, and preANT clades (9,10). Some members of the euANT clade are key regulators of various developmental programs; among these, the PLETHORA (PLT) TFs are particularly important given their roles in regulating embryonic and post-embryonic development (11). The *Arabidopsis thaliana* genome contains six partially redundant *PLT* genes: *PLT1/AINTEGUMENTA LIKE 3* (*AIL3), PLT2/AIL4, PLT3/AIL6*, *PLT4/AIL2*/*BABY BOOM* (*BBM*), *PLT5/AIL5*, and *PLT7/AIL7.* The combined expression of these genes guides developmental programs in the embryo, meristems, and primordia (incipient plant organs) (11–15).

*PLT* gene expression begins during early embryogenesis. *PLT2* and *PLT4/BBM* are expressed from the zygote stage onwards, as they are required for cell division in the zygote and early proembryo (16,17). During the one-cell embryo stage, *PLT2*, *PLT4*, and *PLT7* are expressed in the embryo and *PLT2*, *PLT4*, *PLT7*, and *PLT1* are expressed in the suspensor cell. These genes, along with *PLT5,* are expressed in both the embryo and suspensor cells beginning at the two-cell stage. By the globular stage, all six *PLT* genes are expressed in the embryo. At later stages, the expression patterns of the *PLT* genes shift according to their future expression domains (17). In addition, *PLT2* and *PLT4* redundantly regulate endosperm proliferation and cellularization beginning with the first steps of endosperm formation (16). *PLT4* is also important for somatic embryogenesis, and it was initially characterized as a key regulator of this process in *A. thaliana* (18,19). Post-embryonically, *PLT3*, *PLT5*, and *PLT7* regulate shoot apical meristem (SAM) maintenance and function (20) and redundantly regulate phyllotaxis (12,21,22).

Despite the crucial roles of *PLT* genes in embryo and shoot development, their best understood and most extensively studied function is as master regulators of root development. *PLT1*, *PLT2*, *PLT3*, and *PLT4* have overlapping expression domains in the root apex, where their functions are partially redundant (13,23–25). The primary root of single loss-of-function *plt* mutants exhibits only subtle phenotypes, but *plt1 plt2* double mutants develop short primary and lateral roots with a determinate growth pattern due to exhaustion of the root apical meristem (RAM). This halts root growth, and the plant develops a short, highly branched root system (23). *plt1 plt2 plt3* mutants and segregating *plt1 plt2^+/-^ plt3 plt4/bbm* mutants fail to develop post-embryonic roots (24). Strikingly, the ubiquitous expression of *PLT2* was sufficient to drive the conversion of the *A. thaliana* SAM to a RAM (23,24).

In the root, *PLT* expression and PLT activity peak at the quiescent center and decrease towards the root differentiation zone, with high *PLT* expression in the RAM, moderate expression at the elongation zone, and very low or no expression at the differentiation zone (13,23–25). Furthermore, all *PLT* genes are important for lateral root development: *PLT3, PLT5*, and *PLT*7 are expressed early in incipient lateral root primordia (LRP), (14,21). Most importantly, PLT3, PLT5, and PLT7 activate *PLT1*, *PLT2*, and *PLT4*, which in turn are crucial for RAM establishment in developing lateral roots. Most *plt3 plt7* and *plt3 plt5 plt7* LRPs fail to emerge, and the few LRs that emerge do not grow. Moreover, the *plt3 plt5 plt7* triple mutant develops clusters of LRPs, suggesting that *PLT3, PLT5*, and *PLT7* are also involved in rhizotaxis (14,21).

Despite their prominent roles in plant development, PLT TFs have primarily been studied in *A. thaliana* and (to some extent) a few crop species such as rice (*Oryza sativa*) (26–28) and maize (*Zea mays*) (29); and in the parasitic species *Striga hermonthica* (30). Here we identified putative *PLT* orthologs across Viridiplantae, including representatives of all extant plant clades, and reconstructed their molecular phylogeny. We suggest that the high number of PLT targets could be explained by the presence of intrinsically disordered regions (IDRs) at the N- and C-termini of these TFs. The IDRs might facilitate protein–protein interactions, particularly among TFs, and therefore could help fine-tune target selection in a context-specific manner. Moreover, we constructed a putative gene regulatory network for PLTs at the root meristem based on publicly available transcriptomic data. The inferred network suggests that PLT TFs are key regulators of RNA processing, specifically rRNA metabolism, and ribosome and ribonucleoprotein biogenesis, among other central cellular processes.

## Material and Methods

### Mining of AP2 proteins from Viridiplantae

The *PLT*-like sequences from embryophyte genomes available at Phytozome were retrieved using the PhytoMine tool. For species not included in Phytozome, the Conifer Genome Integrative Explorer (https://congenie.org) was used for *Picea abies* and *Pinus taeda*; GymnoPLAZA (https://bioinformatics.psb.ugent.be/plaza/versions/gymno-plaza/) for *Ginkgo biloba*; SolGenomics Network (https://solgenomics.net) for the Solanaceae species pepper (*Capsicum annuum*), tomato (*Solanum lycopersicum*), eggplant (*S. melongena*), and potato (*S. tuberosum*); and FernBase (https://www.fernbase.org) for *Azolla filiculoides* and *Salvinia cucullata*. AP2-like protein sequences were retrieved from these databases with the built-in BLASTP algorithm using the *A. thaliana* PLT1 amino acid sequence as a query. In addition, the transcriptome of *Isoetes echinospora* (Isoetaceae) (31) was downloaded and the root apex transcriptome of *Pachycereus pringlei* (Cactaceae), *de novo* assembled by the authors (32), was used for analysis. For these transcriptomes, locally run tBLASTn was performed using the PLT1 sequence of *A. thaliana* as a query. For the Cactaceae species *Carnegiea gigantea*, *Lophocereus schottii*, *Stenocereus thurberi*, and *Selenicereus/Hylocereus undatus*, the putative orthologs of *PLT*s identified in the *P. pringlei* transcriptome were searched in the draft genomes (33–35) via locally run BLASTn. AP2-like sequences were retrieved if the BLAST E-value was <1×10^-10^, and the putative ORFs were translated to >100 amino acid sequences. After further filtering, only sequences with two AP2 domains were retained (domain IDs IPR001471 and PF00847 in the InterPro and Pfam databases, respectively).

### Phylogenetic and synteny analyses

A multiple sequence alignment (MSA) of all retrieved sequences was generated with Clustal-Omega (36) and manually curated. The AP2-linker-AP2 region of the MSA was identified using the *A. thaliana* PLT sequences as a reference. This region was extracted and used to identify the best-fitting evolutionary model with ModelTest (37). The AP2-linker-AP2 matrix was used to generate a maximum likelihood tree with RAxML-NG (38) under the JTT-G4 model. The clade containing all PLT proteins from *A. thaliana* was then selected, and the complete protein sequences from this clade were realigned. The PLT MSA was manually curated and used to find the best-fitting evolutionary model. Robustness of the tree topology was assessed using the transfer bootstrap expectation (TBE) method (39). Evolutionary model tests and phylogeny reconstruction were run on the Tepeu server (LCG-UNAM; http://cimi.ccg.unam.mx/es/uati/equipo/5147). A genomic sequence 1 Gb upstream and 1 Gb downstream the *PLT* of interest of a selected species was retrieved in GenBank format. The genomic regions were extracted and tested for collinearity and similarity with Clinker (40), which explores the microsynteny of gene clusters from an “all vs. all” global CDS alignment contained in the GBK files.

### Inference of the PLT structure

The amino acid sequences of PLT-like proteins from each clade were extracted and realigned separately. Disorder probability for individual protein sequences was assessed with ODiNPred (41). Each amino acid was substituted in the MSA matrix by its disorder-propensity value. The MSA matrix was then visualized as a heatmap. Amino acid disorder propensity values were substituted in the MSA using Seqinr, and heatmaps were visualized with Pheatmap in R. The UniProt IDs of the *A. thaliana* PLT proteins were used to retrieve AlphaFold-predicted structures for each PLT. The structural models were visualized with PyMOL (https://pymol.org/2/).

### RNA-seq data analysis

The raw RNA-seq reads from the meristematic, elongation, and differentiation zone (three biological replicates per zone) from *A. thaliana*, *Oryza sativa*, *Zea mays*, soybean (*Glycine max*), cucumber (*Cucumis sativus*), and *Solanum lycopersicum* generated by (42) were downloaded from the GEO Omnibus (accession number: GSE64665). Read quality was assessed with FastQC v0.11.9 (43) and MultiQC v1.14 (44). Read preprocessing was performed with Trimmomatic v0.39 (45) using default parameters. Read mapping was performed by local alignment to the corresponding reference genome with Bowtie2 v2.4.2 (46). The reference genomes in FASTA format and their corresponding GFF3 annotation files were downloaded from the JGI Genome Portal (https://genome.jgi.doe.gov/portal/). The GFF3 files were converted to GFT format with GffRead. Transcript quantification considering overlapping metafeatures (-O) was performed with FeatureCounts v2.0.1 (47) using gene_id and transcripts as features. Genes with low expression counts were discarded; only genes with more than 1 count per million (CPM) mapped reads in at least 3 of the 9 samples per species were retained. Count normalization was performed with DESeq2 (48) implementing a regularized logarithm transformation (rlog) to ensure homoscedasticity and avoid sequencing depth bias. The normalized count matrix was used for principal component analysis (PCA). The Ggplot2 and factoextra packages were used to visualize and extract additional information about the variables and individuals, respectively.

Differentially expressed genes (DEGs) in paired comparisons (meristematic vs. differentiation zone and meristematic vs. elongation zone) were identified using the raw count matrix and the corresponding metadata with DESeq2 using a *lfcThreshold* of log2>1 and an α= 0.05. Genes specifically upregulated in the meristematic zone were defined as those in the intersection of upregulated DEGs in the meristematic zone from each comparison. GO enrichment and Gene Set Enrichment Analysis (GSEA) were performed with clusterProfiler v4.2.2 (49) using the GO annotations for each species provided by Phytozome; *p*-values were adjusted using the Benjamini-Hochberg method. The *p*-and *q*-value thresholds were set to 0.05 and 0.01, respectively.

### Identification of PLT-regulated nodes

The Retrieve Sequence tool from RSAT Plants suite (50) was used to extract the-3000 to-1 region upstream of the transcription start site of each putative PLT target, preventing overlap with neighboring genes (noorf option). The retrieved sequences were used to identify PLT binding sites using position weight matrices (PWM) for the TF binding sites of the PLTs provided by CisBP (51) for *A. thaliana* and their corresponding motifs for the remaining species defined by similarity regression. The RSAT *Matrix-scan* tool was used for pattern matching of the PLT binding sites in the promoter regions with a second-order Markov model, and oligo frequencies were calculated for the corresponding genome assembly for background model estimation.

A list of DEGs in the meristematic zone was obtained from the intersection of three datasets: i) those reported by (13), which correspond to genes with expression changes in seedlings with transiently induced PLT activity; ii) the set of genes specifically upregulated in the meristematic zone, as defined in the previous section, based on data from (42); and iii) genes with PLT-binding sites in their promoter regions.

### Gene Network Analysis

Weighted Gene Correlation Network Analysis (WGCNA) and preservation module analysis were implemented with the WGCNA package v1.72.1 in R (52). The co-expression adjacency matrix with Pearson correlation and module eigengenes for kME calculation were obtained for the rlog normalized count matrix. The dissimilarity topological overlap matrix was used for module detection. The Gaussian mixture model for decomposition per module over the edge weights was used to set a module-specific threshold and discard weak interactions. The *normalmixEM* function from the *mixtools* v2.0 R package (53) was used to perform decomposition of the edge weight distribution into two Gaussian distributions (k=2). Interactions with a probability of >0.5 belonging to the distribution with the lower mean were considered to be weak interactions.

Putative orthologs between species were identified using Bidirectional Best Hit (BBH) criteria in a locally run BLASTp between proteins from each species against *A. thaliana* using BLAST v2.2.29 with default significance thresholds and the R package orthologr v0.4.0 (54). Interspecies network preservation analysis was performed with the table of putative orthologs and the *preservationModule* function of the WGCNA R package, which was run with 200 interactions and biweight midcorrelation to infer co-expression. A network for each root development zone per species was obtained and compared to the *A. thaliana* RAM reference network to assess module preservation (55). Eigengenes for all the datasets were used to calculate kME with the *multiSetMEs* function. The resulting networks were visualized with iGraph (56). For GO enrichment assessment of conserved nodes and interactions across species, the nodes connected by the 100 edges with the lowest variance among all RAM networks were analyzed with the function enrichGO of clusterProfiler v4.2.2 (49), using all the nodes of the reference network as background.

### Single-cell RNA-seq data integration

The inferred PLT GRN was used as a reference interactome for the Single-Cell Imputation and NETwork Construction (SCINET) framework (57) with the dataset GSE152766 (58) and cell-type markers identified using PHYTOMap (Plant hybridization-based targeted observation of gene expression map) (59). Raw single-cell counts were handled according to the ACTIONET framework from the R package ACTIONet v1.0 (57). Normalized logarithmic expression profiles were used to obtain activity scores, impute primary function cell types, calculate signature profiles for each cell type, and transform the data into a lower-dimensional space using the principal components of the transcriptional profiles. The *PLTs* were treated as marker genes to calculate the activity scores of each *PLT* in all cell-type projections. A Gaussian mixture model decomposition per cell-type network was used to set cell type–specific thresholds and discard non-significant interactions in each subset (cell type or root zone) using the normalmixEM function (k=2; 1,000 maximum interactions) from the mixtools v2.0 package (53).

## Results

### Molecular phylogeny of the PLTs

PLT proteins are AP2/EREBP TFs belonging to the AP2 subfamily, as they contain two AP2 domains. We used this feature to identify proteins from the AP2 subfamily in the genomes of embryophyte species by interrogating the Phytozome and other genome databases. We retrieved 1,162 AP2-subfamily protein sequences from 60 plant species belonging to 30 botanical families (Fig 1; S1 Table; S1 Data). These include sequences from the major clades of extant embryophytes: Bryophyta, Lycophyta, Monilophyta, and Spermatophyta; the latter includes *Amborella trichopoda* as a basal species and sister taxon to monocot and eudicot clades. We aligned and manually curated the retrieved sequences. The most suitable evolutionary model for the region comprising the two AP2 domains and the interdomain sequence was the JTT+G4 model (S2 Table). We reconstructed an unrooted maximum likelihood tree (S1 Fig) and identified the clade containing all PLT sequences from *A. thaliana*, as well as the sequences of their sister groups. From this group, we obtained a subset of 316 complete amino acid sequences, which we used to reconstruct the phylogeny of putative PLTs (Fig 2A). These sequences are hereafter referred to as PLT-like sequences (S3 Table).

**Fig 1.**
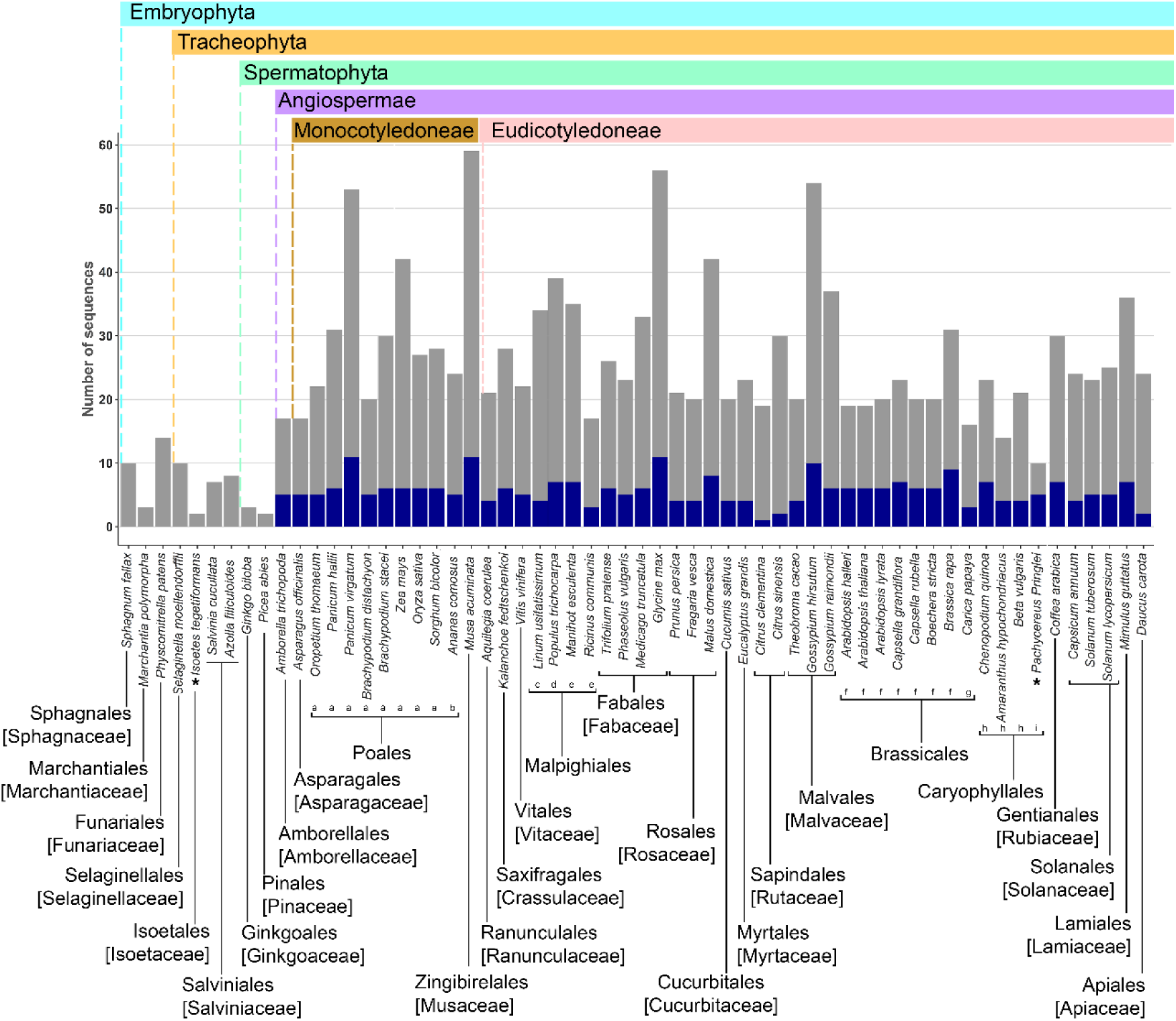
Number of AP2-subfamily genes from the AP2/ERF family in 60 plant species from different taxa. The corresponding protein sequences were used for phylogenetic analysis. Blue bars: *PLETHORA-like* genes; gray bars: *non-PLT AIL* genes. When all species from an order belong to the same family, the family is indicated in brackets below the order. If species from more than one family belong to the same order, the family is indicated with lowercase letters below the species name as follows: a. Poaceae, b. Bromeliaceae (Poales); c. Linaceae, d. Salicaceae, e. Euphorbiaceae (Malpighiales); f. Brassicaceae, g. Caricaceae (Brassicales); h. Amaranthaceae, i. Cactaceae (Caryophyllales). *The transcriptome of *Isoetes echinospora* leaves, corms, and rootlets and the root apex transcriptome of *Pachycereus pringlei* were used to identify genes encoding proteins with two AP2 domains; therefore, more paralogs could exist in their genomes.

**Fig 2.**
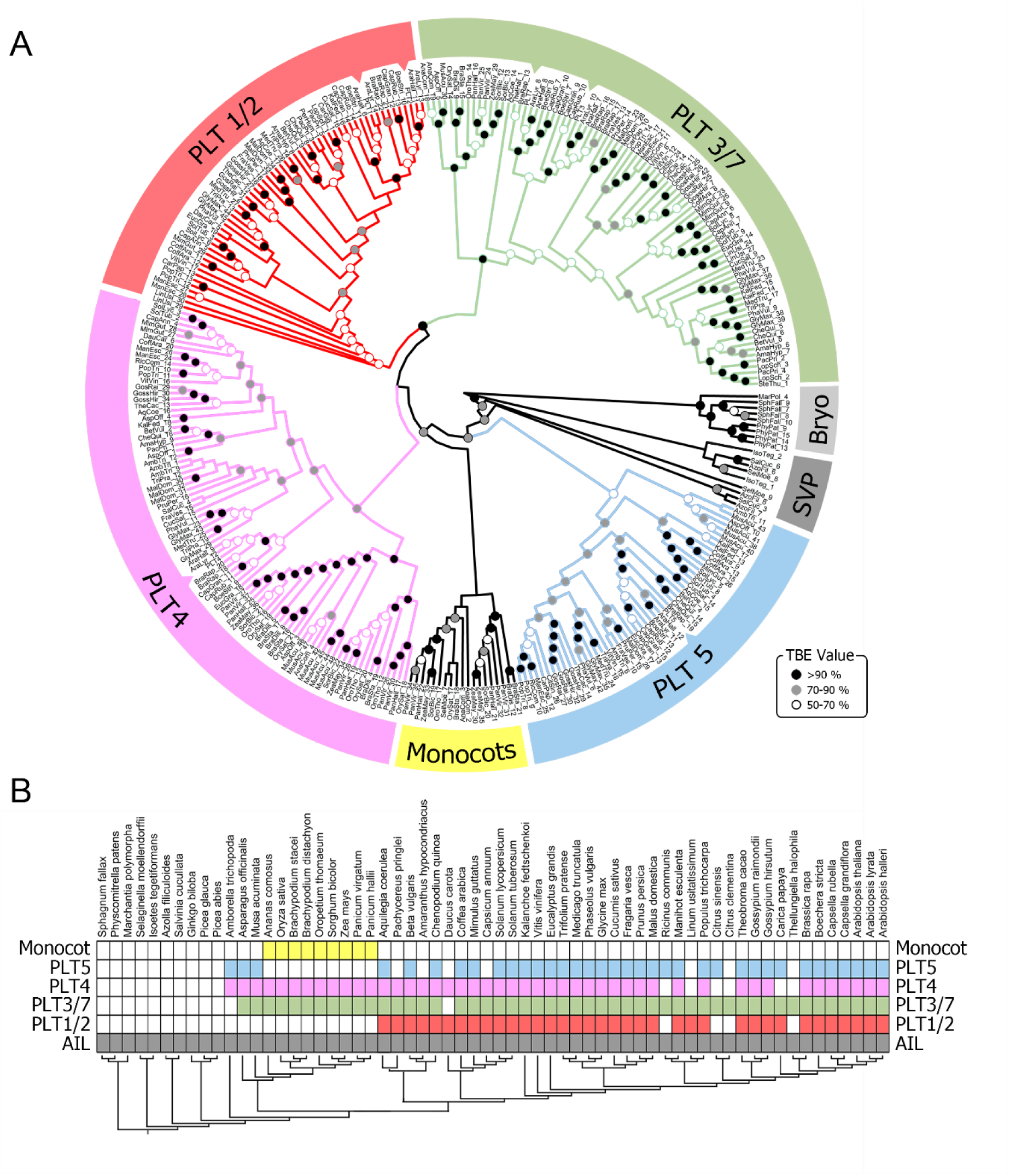
Maximum likelihood phylogenetic tree of PLETHORA-like proteins and distribution of AP2-subfamily proteins of each species in the clades. **A.** Phylogeny of the selected dataset of deduced PLT-like proteins, as well as AIL proteins from non-angiosperm embryophytes: Bryophyta (Bryo) and seedless vascular plants (SVP); see S1 Fig. for the phylogeny of the complete dataset of AP2-subfamily proteins used in this study. The gene identifiers for the sequences are listed in S1 Table. Branch support values were calculated using the transfer bootstrap expectation (TBE) algorithm, with 1,000 replicates. TBE values are indicated by circles on the branches. Four PLT clades are named according to their putative *A. thaliana* orthologs, indicated by dents in the respective ribbon. The PLT3/7 clade is shown in green, PLT1/2 in red, PLT4 in pink, and PLT5, which is sister to the other PLT clades, in blue. **B.** Placement of the proteins used for the phylogeny within recovered clades. Colored rectangles: the species contains at least one PLT-like protein from the respective clade; white rectangles: the species does not contain PLT-like proteins from the respective clade.

We identified putative AP2/EREBP proteins in the genomes of all analyzed plant species, which supports the previous finding that these proteins have been present since green algae, the sister clade of Embryophyta (3,10). We reconstructed a maximum likelihood tree using the complete sequences of the PLT-like proteins and calculated branch support with the TBE algorithm, with 1,000 replicates (39). The resulting tree topology is divided into five well-supported clades. Four of these clades were named based on the *A. thaliana* PLTs they contain: (*i*) PLT1 and PLT2, (*ii*) PLT3 and PLT7 (*iii*) PLT4, and (*iv*) PLT5 (Fig 2A). The PLT4 (BBM) clade was resolved as sister to the PLT1-PLT2 and PLT3-PLT7 clade, while PLT5 was resolved as the sister clade to all other PLT clades (Fig 2A). The tree topology supports the loss of PLT1-PLT2 in monocots, which was possibly compensated for by the emergence of the fifth PLT clade, which is monocot-specific (Fig 2A) (8,10).

### The *PLT* clade arose before the divergence of angiosperms

The newly generated tree topology is mostly congruent with previous phylogenetic reconstructions, in which PLT1 is grouped with PLT2, and PLT3 is grouped with PLT7. To assess the duplication of genomic regions, either from segmental or whole genome duplication events, we extracted the nucleotide sequences near the *A. thaliana PLT* loci and evaluated gene cluster similarity. Very strong microsyntenic signals were detected for the genomic regions of *PLT3* and *PLT7* (both on chromosome 5), and strong signals were detected between *PLT1* and *PLT2* (located on chromosomes 3 and 1, respectively) (Fig 3A). No syntenic relationship of *PLT4* or *PLT5* with any other *A. thaliana* paralog was observed.

**Fig 3.**
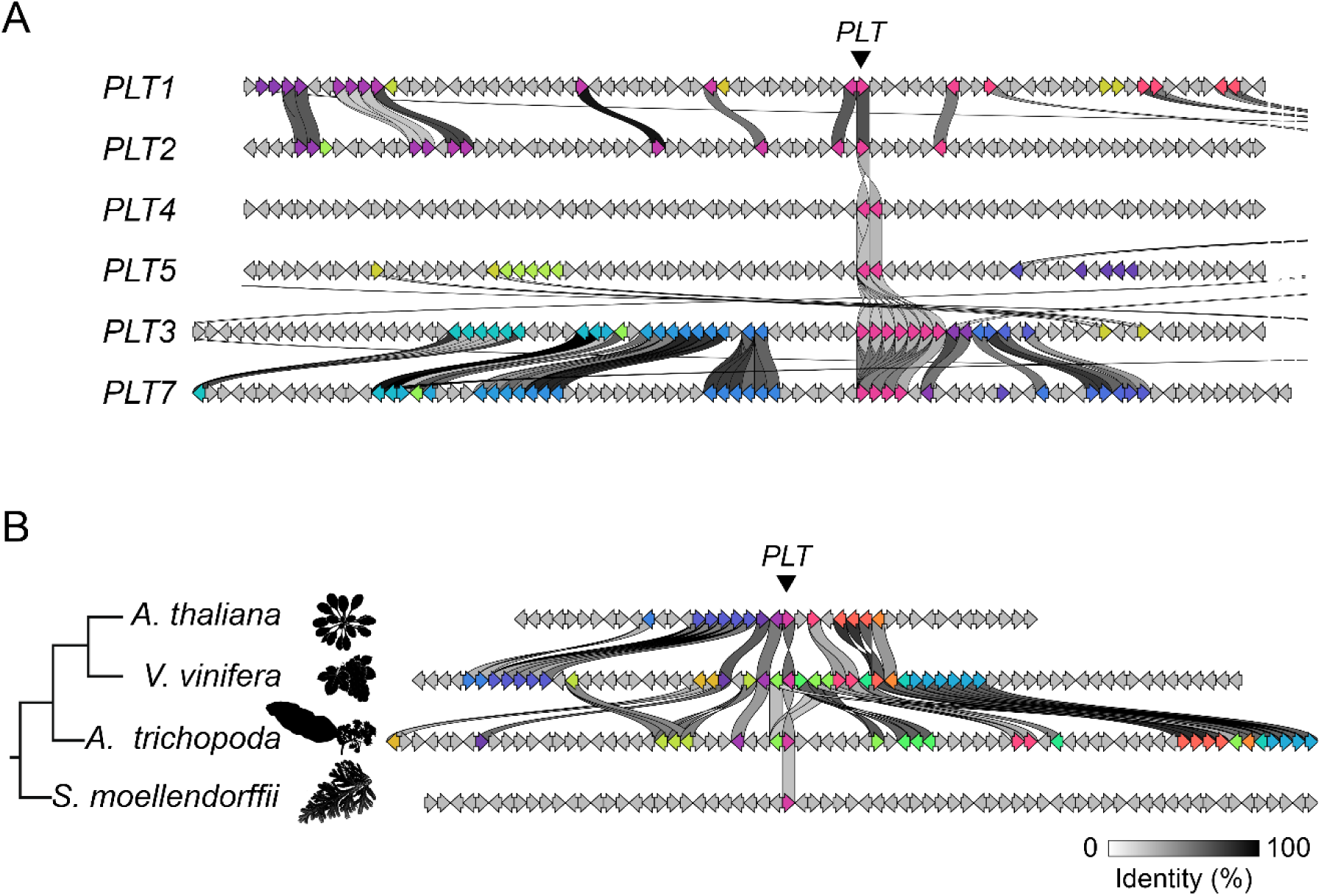
Analysis of microsynteny of the six *PLETHORA* genomic regions in *Arabidopsis thaliana* (A) and the *PLT1-like* region in four plant species (B). The Clinker tool was used to analyze the genomic sequences 1 Gb upstream and 1 Gb downstream of the *PLT* of interest. **A**. The *PLT3* and *PLT7* genomic regions show highly conserved gene orders, supporting the notion that they originated from a recent duplication. The *PLT1* and *PLT2* regions also show significantly conserved gene order, while neither the *PLT4* nor *PLT5* region shows conserved synteny with other *PLT* regions. **B**. The eudicots *Arabidopsis thaliana* and *Vitis vinifera*, as well as *V. vinifera* and *Amborella trichopoda* (the sister group to other angiosperms), show strong signs of conserved gene order, whereas the seedless vascular plant *S. moellendorfiii* does not show such signs with any of the three other species.

We then asked to what extent this syntenic conservation could still be observed among basal plants. We used chromosomal regions centered on the *PLT1-like* loci of selected species, including *Selaginella moellendorffii*, *Amborella trichopoda*, grapevine (*Vitis vinifera*), and *A. thaliana*. We selected these species based on their divergence times and placement in the Embryophyte lineage. Strong syntenic conservation was observed between the chromosomal regions of *A. thaliana* and *V. vinifera*, the latter as a representative of species diverging at the base of the Pentapetalae (core eudicots) clade (Fig 3b). Furthermore, the syntenic conservation of this region was still observed when we compared *A. thaliana* and *Amborella trichopoda*; therefore, this result traces the origin of PLTs back to the base of the angiosperm lineage. However, synteny conservation was not evident between *A. thaliana* and *S. moellendorffii*, a seedless vascular plant (Figs 1 and 2).

### PLT proteins likely contain intrinsically disordered regions

Given that many plant TFs are enriched with IDRs (60,61), we looked for likely disordered regions within stretches of the PLT amino acid sequences (Fig 4A, S2-S5 Figs). This analysis revealed a core structured region, which corresponds to DNA binding domains (the AP2-interdomain-AP2 region), and likely disordered regions towards the N- and C-termini of the proteins. Indeed, the AlphaFold-predicted structures for all *A. thaliana* PLT proteins suggest that the N- and C-termini might be unstructured (Fig 4B, S2–S5 Figs), as indicated by the low confidence values at these regions (S6 Fig).

**Fig 4.**
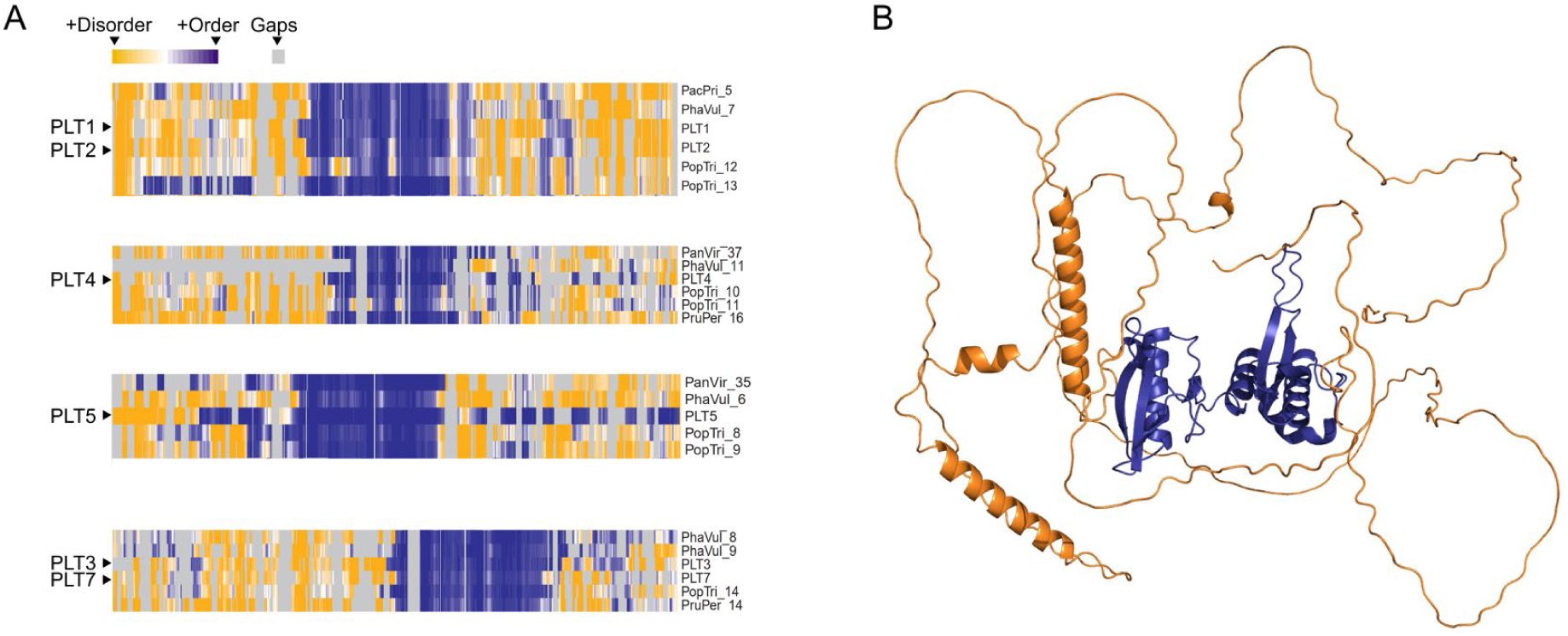
PLETHORA transcription factors are predicted to contain intrinsically disordered regions. **A.** Intrinsic disorder in six PLETHORA proteins of monocot and eudicot species predicted by ODiNPred. *A. thaliana* proteins are listed without the species acronym; *Pacpri*: *Pachycereus pringlei*; *Phavul*: *Phaseolus vulgaris*; *Panvir*: *Panicum virgatum*; *Poptri*: *Populus trichocarpa*; *Pruper*: *Prunus persica*. Intrinsic disorder predictions for all PLT-like proteins in each clade are provided in S2–S5 Figs. **B.** AlphaFold-predicted structure of *Arabidopsis thaliana* PLT1, a representative structure. The predicted structures of six *A. thaliana* PLTs are shown in S6 Fig.

### Gene Regulatory Network

To infer a Gene Regulatory Network (GRN) mediated by the PLT TFs, we used the transcriptomes of developmental zones of the primary root in six angiosperm species generated by Huang and Schiefelbein (42). We first identified DEGs that were upregulated in the meristematic zone but not in the elongation or differentiation zone, finding 5,086 genes that were specifically upregulated in the meristematic zone of *A. thaliana* (S4 Table, S7 Fig). To compare the GRNs between species (see below), we also conducted this analysis for all other species, i.e., *C. sativus*, *G. max*, *S. lycopersicum*, *O. sativa*, and *Z. mays,* and identified 3,854, 5,456, 5789, 3,242, and 5,552 DEGs, respectively, in the meristematic zone compared to the elongation and differentiation zones (S4 Table). To identify genes that exhibit changes in expression when *PLT* expression is altered, we used the *A. thaliana* transcriptomes of seedlings with dexamethasone-inducible activity of each PLT TF (13). We identified 638 DEGs in the RAM with altered expression in seedlings following the induction of PLT activity. To discriminate between putative direct and indirect targets of PLTs, we scanned the 3 Kb region upstream of the transcription start site of each putative target gene and retained only sequences that included a PLT binding motif. This analysis further reduced the number of putative PLT targets to 534 genes (Fig 5A).

**Fig 5.**
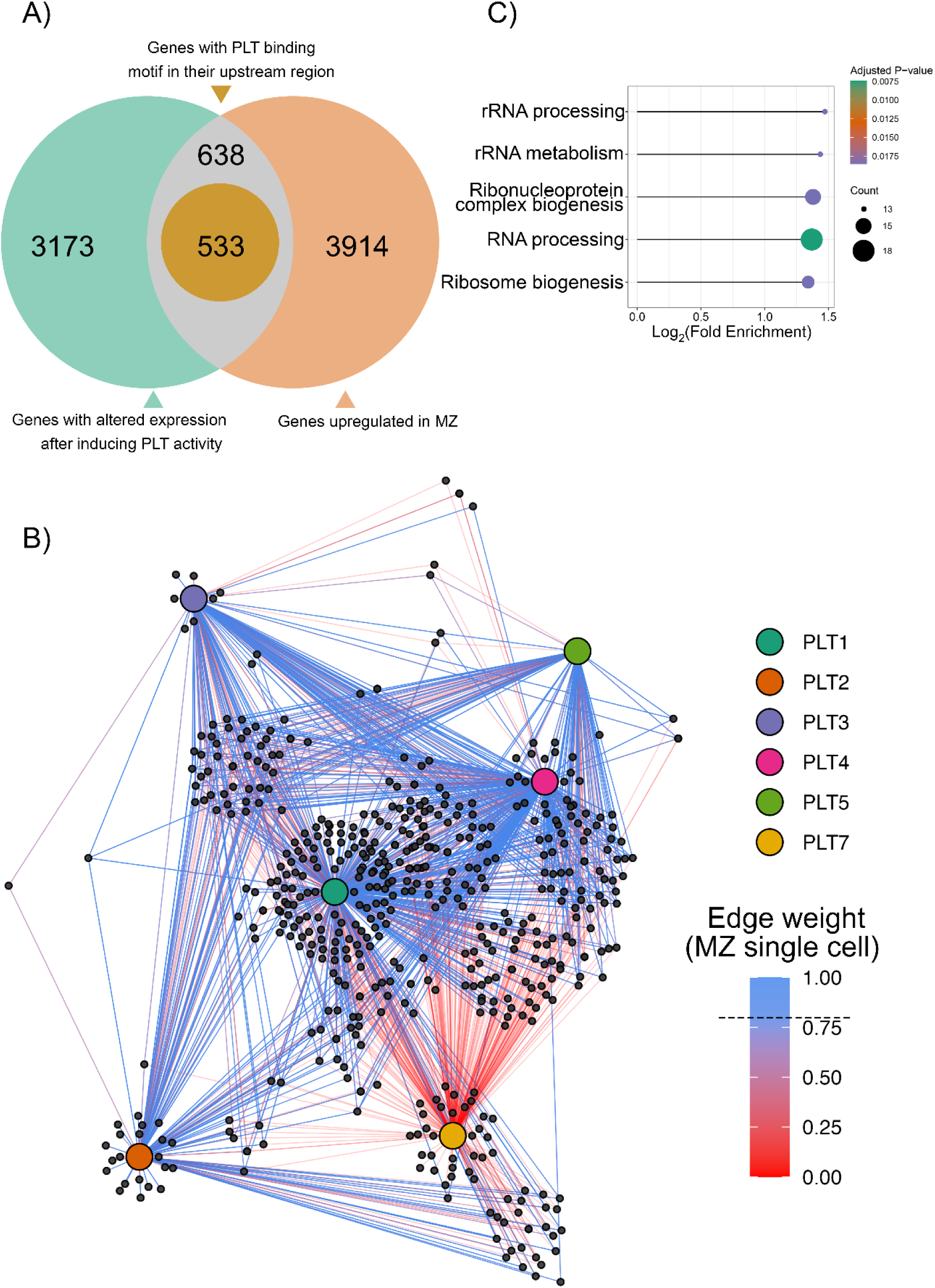
Gene regulatory network of PLETHORA transcription factors in the *Arabidopsis thaliana* root apical meristem. **A.** Venn diagram of the number of differentially expressed PLT target genes in the RAM that possess a PLT binding motif upstream of the transcription start site. **B.** Undirected regulatory network with 1,444 putative interactions, including the 6 PLTs and 533 putative targets. Edge weight in the network derived from single-cell RNA sequencing is shown by the red-blue gradient. The threshold to discriminate significant (>0.79) from non-significant (<0.79) interactions (dashed line) was selected by Gaussian mixture model decomposition analysis (see S10 Fig). **C.** GO enrichment analysis of biological processes performed over the nodes with the lowest interspecies variance in the PLT network. The most over-represented GO terms were ribosome and ribonucleprotein complex biogenesis and RNA processing.

We constructed an undirected network with 1,444 putative interactions, or edges (S5 and S6 Tables), using 539 nodes (6 PLTs + 533 putative targets) and visualized the network with iGraph (Fig 5B). As an independent approach to assess the GRN mediated by PLTs, we implemented Weighted Gene Correlation Network Analysis (WGCNA), an unsupervised algorithm that leverages co-expression profiles to infer regulatory modules (52). Using the complete and unfiltered *A. thaliana* root RNA-seq data (42) generated from samples of the meristematic, elongation, and differentiation zones of the primary root, we obtained 39 modules (S8 Fig). All *PLTs*, except *PLT7*, were included in the largest (turquoise) module (S8A,B Fig), which contained 505 of the 539 nodes included in the manually curated network. To assess whether the network is associated with specific cellular processes, we analyzed the enrichment of GO biological processes represented by the network nodes. The biological processes enriched in the turquoise module of the WGCNA were related to RNA metabolic processes, including rRNA metabolism and processing, ribosome and ribonucleoprotein biogenesis, as well as mRNA and non-coding RNA metabolism, splicing, and modification (S8C Fig).

Given the central role of rRNA in cellular processes, we asked whether the PLT-mediated network is conserved across angiosperms. We generated a WGCNA network for *G. max*, *C. sativus*, *S. lycopersicum*, *Z. mays*, and *O. sativa* as previously described using the root RNA-seq data from the same source (42). We calculated density and connectivity scores for each network and combined them as Z-summary statistics by comparing them to the *A. thaliana* reference network. The Z-summary value for all species was >2, providing evidence for conservation, and >10 in the meristematic zone for *C. sativus*, *S. lycopersicum*, and *O. sativa*, providing strong evidence for conservation of the GRN in these species (55) (S9 Fig). We then extracted the gene IDs of the network nodes from each species, retrieved their amino acid sequences, and used the sequences to perform BBH BLASTp analysis. This allowed us to calculate the identity and similarity of the individual nodes across species (S10 Fig). Our results suggest that most nodes in the network are strongly conserved in monocots and eudicots, with a median identity and similarity percentage >75% (S10 Fig). By performing an enrichment analysis on the network nodes with less variance across species, in line with our previous result, we found that the most over-represented categories were related to ribosome biogenesis, including rRNA processing, rRNA metabolism, and ribonucleoprotein complex biogenesis (Fig 5C).

Both the manually curated and WGCNA networks were derived from bulk RNA-seq data, which might obscure fine-tuned regulation in a cell type–specific context. To address this issue, we integrated single-cell transcriptomic data (58,59) to weight interactions according to gene activity values in particular cell types (57). For this analysis, we calculated gene activity, i.e., a scaled and normalized distribution of gene expression values comparable across genes and cell types (S11A Fig). We then used the gene activity values to weight the likelihood of gene–gene transcriptional interactions, i.e., to assign occurrence likelihood scores to the edges of the network, using Gaussian mixture model decomposition (S11B Fig). From this analysis, we identified putative interactions of individual PLTs and observed distinct interactions mediated by PLTs across root development zones (Fig 5B, S11B Fig).

## Discussion

PLT proteins are master regulators of several developmental programs in plants (11). However, reconstructing their phylogeny and uncovering their genetic regulatory mechanisms are challenging due to their high similarity, overlapping expression domains, and partial functional redundancy (13,23,24). Although a few phylogenomic studies of AP2/EREBP proteins were recently reported, these studies were performed for the whole AP2/EREBP superfamily using large datasets but focusing only on angiosperms (8) or using wide-spanning datasets but with low-confidence resolution of the PLT clade (10).

Here, to fill these gaps, we present a molecular phylogeny of PLT TFs from representative species of all major taxa of extant plant species (Fig 1 and 2). AP2 proteins originated in the common ancestor of Viridiplantae (3), indicating that this plant-specific TF family arose prior to the colonization of land, and therefore, that PLT TFs might be present in early divergent embryophyte clades. Our analysis suggests the existence of ANT and AIL proteins but not PLT-like proteins in Bryophyta, Lycophyta, and Monilophyta. Although the available genomes of early divergent species are scarce, especially for Lycophyta and Monilophyta, this observation suggests that PLT proteins might have emerged by neofunctionalization of AIL proteins prior to the emergence of seed plants (Fig 1). We were unable to retrieve any PLT-like sequences from the proteomes of the gymnosperm clade, even though a putative ortholog of *PLT4* was previously reported for *Picea abies* (10). This can be explained by the assembly status of gymnosperm genomes, which are mostly fragmented and in draft versions, and the high thresholds used here for E-value and length of the encoded protein, which could overlook putative PLT-like sequences in gymnosperms.

Given that the reconstructed phylogenetic tree includes sequences from evolutionarily distant species, the classic Felsenstein bootstrap method to test tree topology robustness was deemed inappropriate, as the bootstrap support values decrease abruptly with increasing taxon sample size and depth (39,62–64). As an alternative, we used the TBE algorithm, which is more suitable for reconstructing single-gene phylogenies based on large datasets from distantly related species (39,64). The resulting embryophyte PLT phylogeny (Fig 2) is in line with the general tree topology observed for *A. thaliana* PLT proteins (8) and suggests that *PLT1* and *PLT2*, as well as *PLT3* and *PLT7*, likely arose from recent gene duplication events. This hypothesis is supported by the discovery that whole genome duplication events are widespread the evolutionary history of Viridiplantae (65) and by the results of synteny analysis, which revealed conserved syntenic blocks between *A. thaliana* (Brassicales) and *V. vinifera*, as well as between *V. vinifera* and *Amborella trichopoda* (Amborellales) (Fig 3). The results of synteny analysis suggest that the ancestral sequence of PLT1/PLT2 existed in the ancestor of angiosperm species. However, no sequences from the PLT-like clade were identified in *Selaginella moellendorffii* (Selaginellales). The lack of conserved synteny with this seedless vascular plant further suggests that PLTs were not present in early divergent Tracheophyta (Figs 1-3). These observations support the hypothesis that PLT TFs arose by neo-functionalization from an AIL ancestor. However, not enough genomic information is available for Gymnosperm species to test whether PLT proteins originated during the divergence of the Spermatophyta (gymnosperms + angiosperms) or Anthophyta (angiosperms) clade, a question that requires further exploration.

The separation of the PLT1-PLT2 and the PLT3-PLT7 clades points to functional divergence of the PLT proteins, which is supported by the roles of PLT1 and PLT2 as regulators of RAM maintenance and the recruitment of PLT3, PLT5, and PLT7 as regulators of lateral root primordium organogenesis, phyllotaxis, and rhizotaxis (12,14,21).

Our phylogenetic analysis supports the hypothesis that monocot species lost the *PLT1* and *PLT2* genes (Fig 2). In addition, we were unable to identify *PLT5* in some monocot species. The complete loss of *PLT1* and *PLT2* in the monocots surveyed is intriguing, given the prominent role of these TFs in primary root growth. The phylogeny also points to the existence of a monocot-specific PLT clade, which might have compensated for the loss of *PLT1* and *PLT2* orthologs. The loss of *PLT1* and *PLT2* in monocots and the emergence of the monocot-specific clade were previously proposed to be related to the evolution of the fibrous root system architecture of monocots, which is distinct from the tap root system of most eudicot species (8). Our results strengthen this hypothesis. Notably, we identified two *PLT1-PLT2* orthologs, two *PLT3-PLT*7 orthologs, and one *PLT4* ortholog in the *P. pringlei* root apex transcriptome and in the preliminary genomes of five Cactaceae species (S1 Data). The expression of these *PLT* genes in the cactus root apex suggests that the development of the fibrous root system is regulated by distinct mechanisms in monocots vs. Cactaceae, which exhibits determinate growth of primary and lateral roots (66,67).

One of the most challenging features of studying PLT-mediated regulation is the partial overlap of PLT expression domains and their functional redundancy (13,24,25). The results from protein disorder analysis suggest that PLT proteins might contain IDRs towards their N- and C-termini (Fig 4). This is in agreement with the previous finding that eukaryotic TFs, especially in plants, contain a high proportion of IDRs (61). IDRs tend to be located in the activation domains of TFs, where they might increase the number of interactors a TF can recognize, and therefore, the number of putative targets they can regulate (61,68). Some members of the AP2/ERF superfamily contain long IDRs (69). Moreover, Burkart *et al*. (2022) showed that PLT3 interacts with the TF WUSCHEL RELATED HOMEOBOX 5 (WOX5) to control columella stem cell specification (70), and Shimotohno *et al.* (2018) demonstrated that PLT physically interacts with TCP family TFs (71). Interestingly, TCP family members (72) and WOX5 are thought to contain IDRs. In addition, the PLT3–WOX5 association depends on the presence of prion-like domains in the PLT protein (70). Together, these observations support the hypothesis that PLT proteins interact with their protein partners via IDRs to modulate the recognition of binding sites in the regulatory regions of their targets. This mechanism could account for both the high number of PLT targets and their differential regulation along the root axis to orchestrate root zonation, i.e., cell proliferation, elongation, and differentiation.

To further explore the gene regulatory network controlled by PLT TFs, we built a manually curated GRN and an independently inferred network via WGCNA, an unsupervised algorithm. Strikingly, 94% (505 of 539) of the nodes were included in both the WGCNA and manually curated network. According to this GRN, the main processes regulated by PLTs at the RAM are related to rRNA processing and metabolism (Fig 5). To assess whether the inferred network is biologically informative, we looked for nodes in the network that represent genes previously characterized as part of the PLT regulatory network. We identified ROOT GROWTH FACTOR 1 (RGF1) and RGF3, which are small regulatory peptides that induce a signaling cascade leading to *PLT1* and *PLT2* expression (73). We also identified *A. thaliana* RGF RECEPTOR 1 (RGFR1) and RGFR2 in the PLT GRN network. These RGFRs redundantly recognize RGFs and control RAM size, thereby affecting root length (74). The RGFs and RGFRs act together for the correct specification of the PLT gradient underlying root zonation.

In addition to identifying known regulators of PLT expression, we also identified new putative nodes of the PLT regulatory network, some of which, when their function is abolished, lead to shorter roots, a smaller RAM, or even RAM exhaustion. For example, *RETARDED ROOT GROWTH* (*RRG,* AT1G69380) was identified as a putative PLT target. RRG is a nucleus-encoded mitochondrial protein that promotes cell proliferation and represses cell elongation in the RAM of *A. thaliana*; the loss of function of RRG severely impairs root growth (75). *POPCORN* (*PCN*, AT4G07410), encoding a WD-40 protein, was also identified as a putative PLT target. *pcn* loss-of-function mutants develop short roots with a small RAM and differentiated cells close to the root apex (76), similar to the phenotypes of the *plt1 plt2* double mutant (24). The PCN homolog in yeast, U3 SMALL NUCLEOLAR RNA-ASSOCIATED PROTEIN 4 (UTP4, https://swisspalm.org/proteins/Q969X6), is a snoU3- ribonucleoprotein implicated in ribosome biogenesis (77). This is in line with the finding that the GO term ribosome biogenesis was overrepresented among PLT-regulated processes (Fig 5C).

*NUCLEOSTAMIN-LIKE1* (*NSN1*, AT3G07050), encoding a nucleolar GTP-binding protein, was also identified as a PLT target. NSN1 is required for ribosomal assembly; its loss of function in *A. thaliana* leads to defects in embryo development (78) and post-embryonically to the development of short primary roots with a smaller RAM and a differentiation zone closer to the root apex (79). The *NSN1* homolog in animals was first recognized as a stem cell marker (80) and later as a regulator of stem cell proliferation and differentiation (81). NSN1 is thought to regulate organ growth by modulating ribosome biogenesis in *A. thaliana* (82). *SMALL ORGAN 4* (*SMO4*; AT2G40430), encoding a nucleolar protein, was also identified among putative PLT targets involved in regulating RAM size. *SMO4* is highly expressed in the RAM and other plant organs undergoing cell proliferation (83). The loss of function of *SMO4* leads to a slightly shorter RAM due to an impaired cell cycle. SMO4 is a ribosome biogenesis factor involved in 5.8S and 18S rRNA biogenesis (84). Although the macroscopic phenotypes of the *smo4* loss-of-function mutant are subtle, its phenotype at the cytological level is striking, with nucleolar hypertrophy and disorganization, reflecting the role of SMO4 in ribosome biogenesis (84).

Among putative PLT-regulated genes, we also identified genes encoding the nucleolar proteins REBELOTE (RBL, AT3G55510), SLOW WALKER 2 (SWA2, AT3G55510), NUCLEOLAR COMPLEX ASSOCIATED 2 (NOC2, AT2G18220), and AtNOC3 (AT1G79150). RBL interacts with SWA2, NOC2, and AtNOC3 (85), which together control cell proliferation by regulating ribosome biogenesis (85,86). We also identified *ARABIDOPSIS HOMOLOG OF YEAST RRS1* (ARRS1, AT2G37990), encoding a protein involved in rRNA processing and ribosome biogenesis (87), and a gene encoding putative *A. thaliana* TAF RNA Polymerase I subunit A (AT1G53200) among PLT target genes. Overall, the novel nodes present in the GRN built for PLTs at the RAM support the hypothesis that PLT TFs regulate conserved developmental processes in plants, including rRNA metabolism and ribosome biogenesis. The paramount importance of these processes might account for the evolutionary stability of PLT-mediated regulation, as highlighted by the Z-summary score of the network reconstructed for some eudicot (*C. sativus*, *G. max*, *S. lycopersicum*) and monocot (*O. sativa* and *Z. mays*) species (S9 Fig).

## Conclusion

The gradient of *PLT* gene expression and protein activity provides the root with a mechanism to dictate different cell state outputs along the root axis. PLT proteins in *A. thaliana* primary roots act in a dose-dependent manner, translating their activity into two distinct cellular responses: cell proliferation in the RAM, and a gradient of cell differentiation shootward. We suggest that the complexity of the GRN mediated by PLT TFs could be explained, at least in part, by the IDRs in the N- and C-termini of PLT proteins, which fine-tune target selection along the root axis or across cell types. In addition, we propose that PLTs regulate rRNA metabolism and ribosome biogenesis, providing new directions for elucidating the molecular mechanisms behind the roles of PLT TFs as master regulators of root development.

## Supporting information

### Supplemental figures

**S1 Fig.**
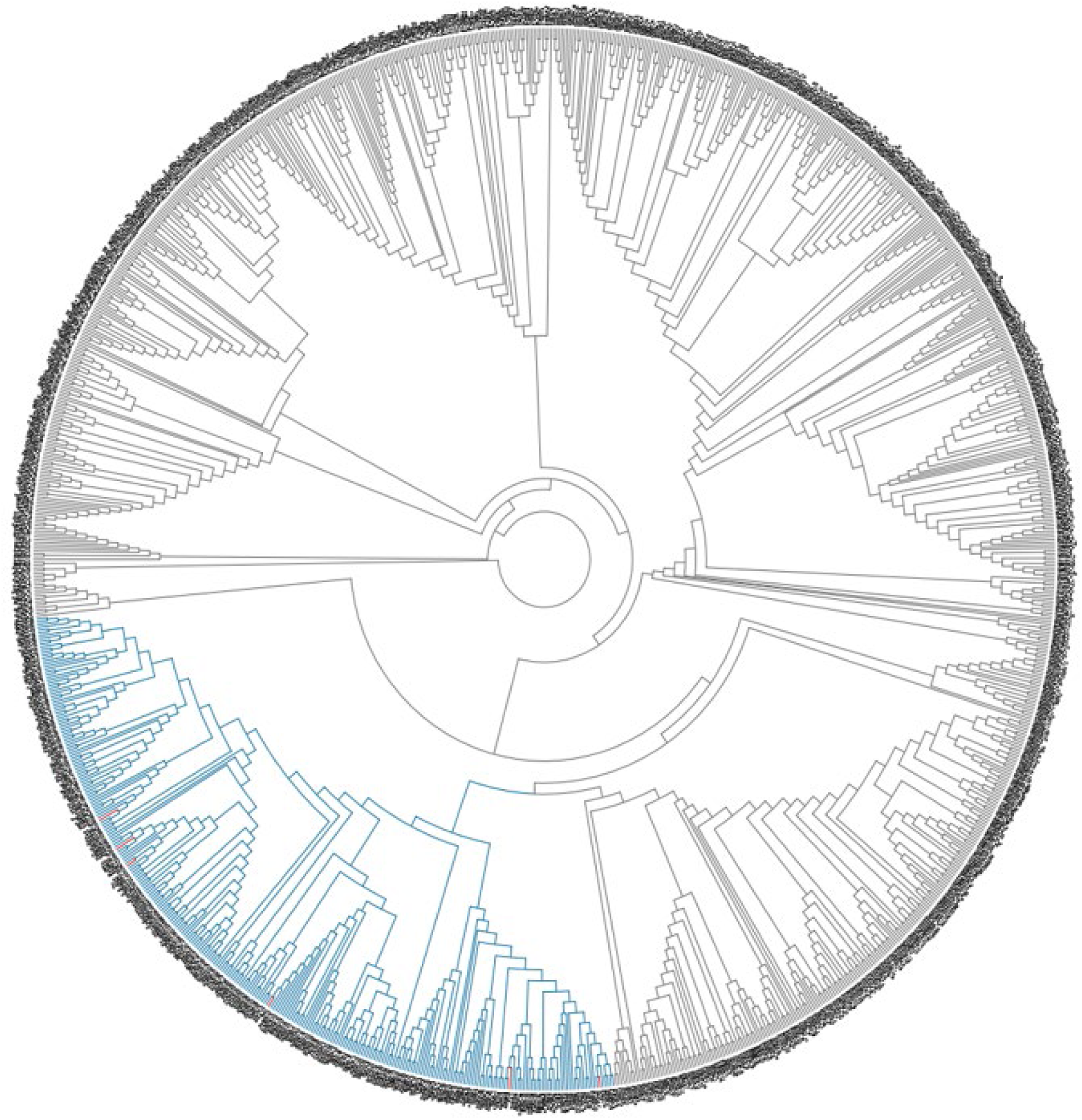
Neighbor-joining phylogenetic tree of AP2 protein sequences in Viridiplantae. The sequences were retrieved from Viridiplantae genomes as described in the Material and Methods. Only protein sequences that included two consecutive AP2 domains separated by an interdomain linker were used for analysis. All PLT sequences from *Arabidopsis thaliana* (red branches) are included in a single clade (shown in blue). The complete protein sequences included in this clade were extracted and realigned (referred to as PLT-like sequences). Information about the sequences used to reconstruct this phylogeny, including database identifiers, is provided in S1 and S3 Tables. The “*FigS1_Tree.nwk”* file is uploaded as a supplemental material.

**S2 Fig.**
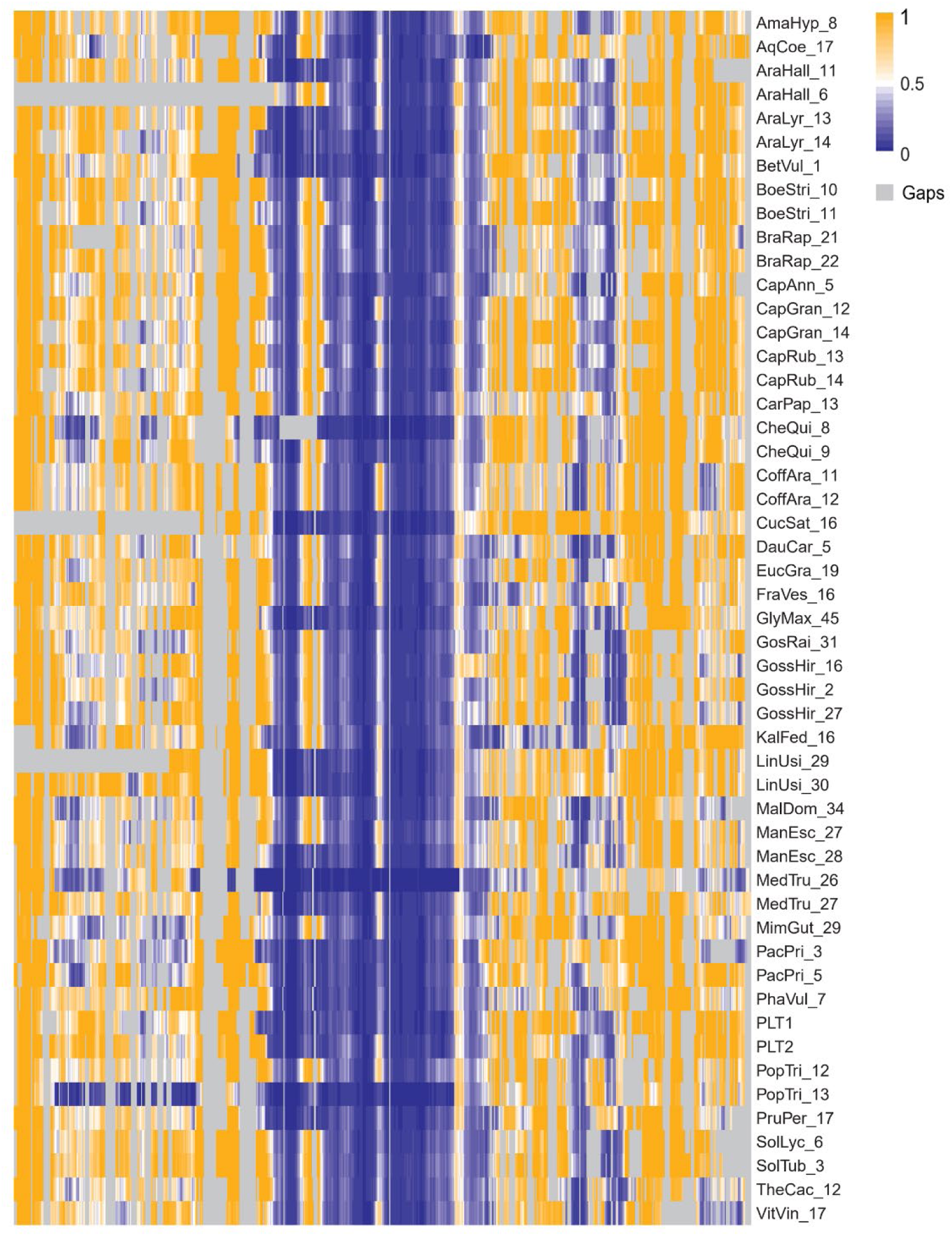
Disorder propensity of putative orthologs of *Arabidopsis thaliana* PLETHORA1 and PLETHORA2. Disorder propensity was analyzed for each PLT sequence of the PLT1-PLT2 clade (Fig 2). Each amino acid in the multiple sequence alignment was substituted by its disorder propensity value. The resulting matrix was visualized as a heatmap. Note that this analysis suggests that the AP2-linker-AP2 region, which corresponds to the DNA binding domains is structured (blue), in line with a crystal structure obtained for a single AP2 domain of TEM1 TF (7ET4) (88). For complete species names, see S3 Table.

**S3 Fig.**
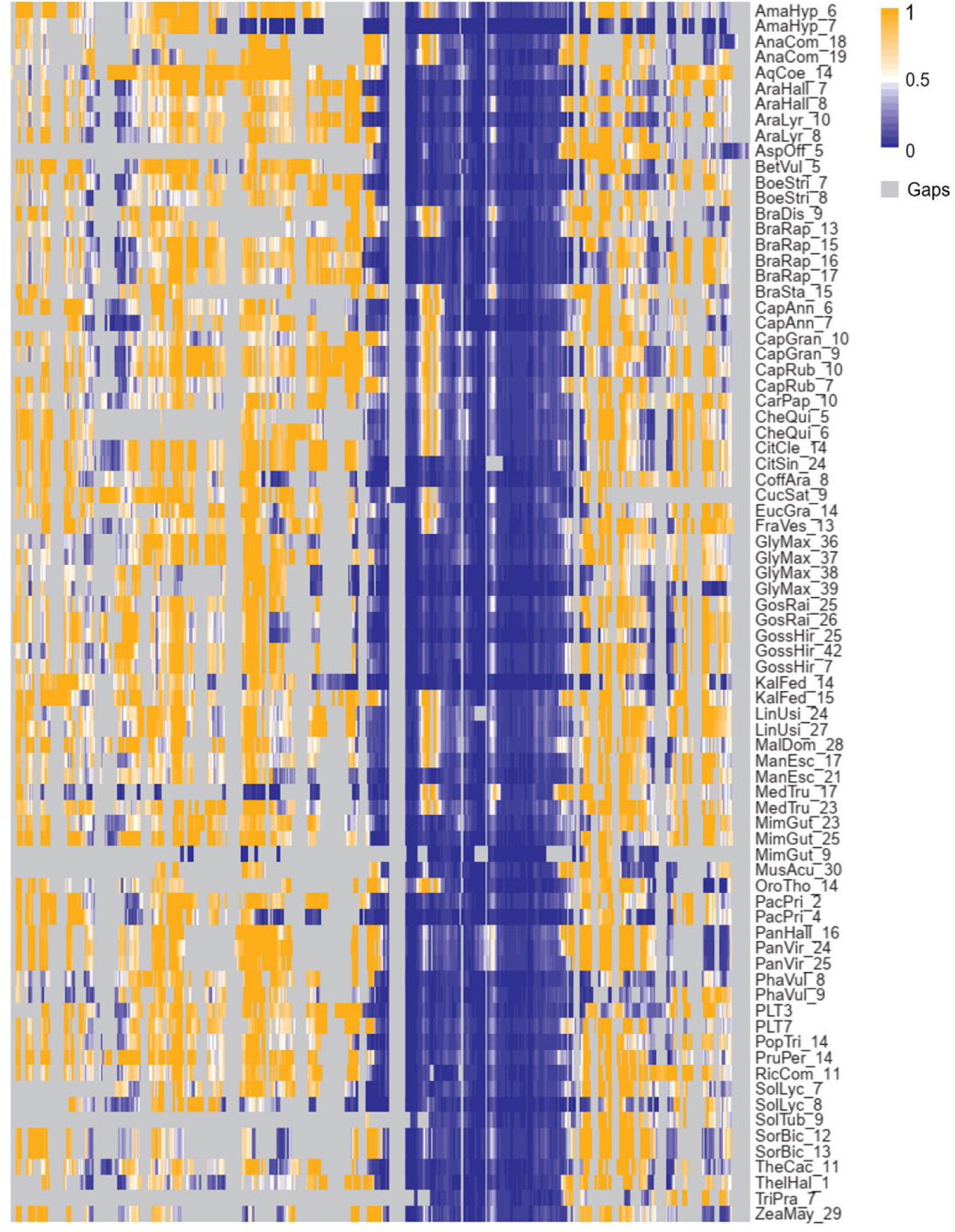
Disorder propensity of putative orthologs of *Arabidopsis thaliana* PLETHORA3 and PLETHORA7. Disorder propensity was analyzed for each PLT sequence of the PLT3-PLT7 clade (Fig 2). Each amino acid in the multiple sequence alignment was substituted by its disorder propensity value. The resulting matrix was visualized as a heatmap. Note that this analysis suggests that the AP2-linker-AP2 region, which corresponds to the DNA binding domains, is structured (blue), in line with a crystal structure obtained for a single AP2 domain of TEM1 TF (7ET4) (88). For complete species names, see S3 Table.

**S4 Fig.**
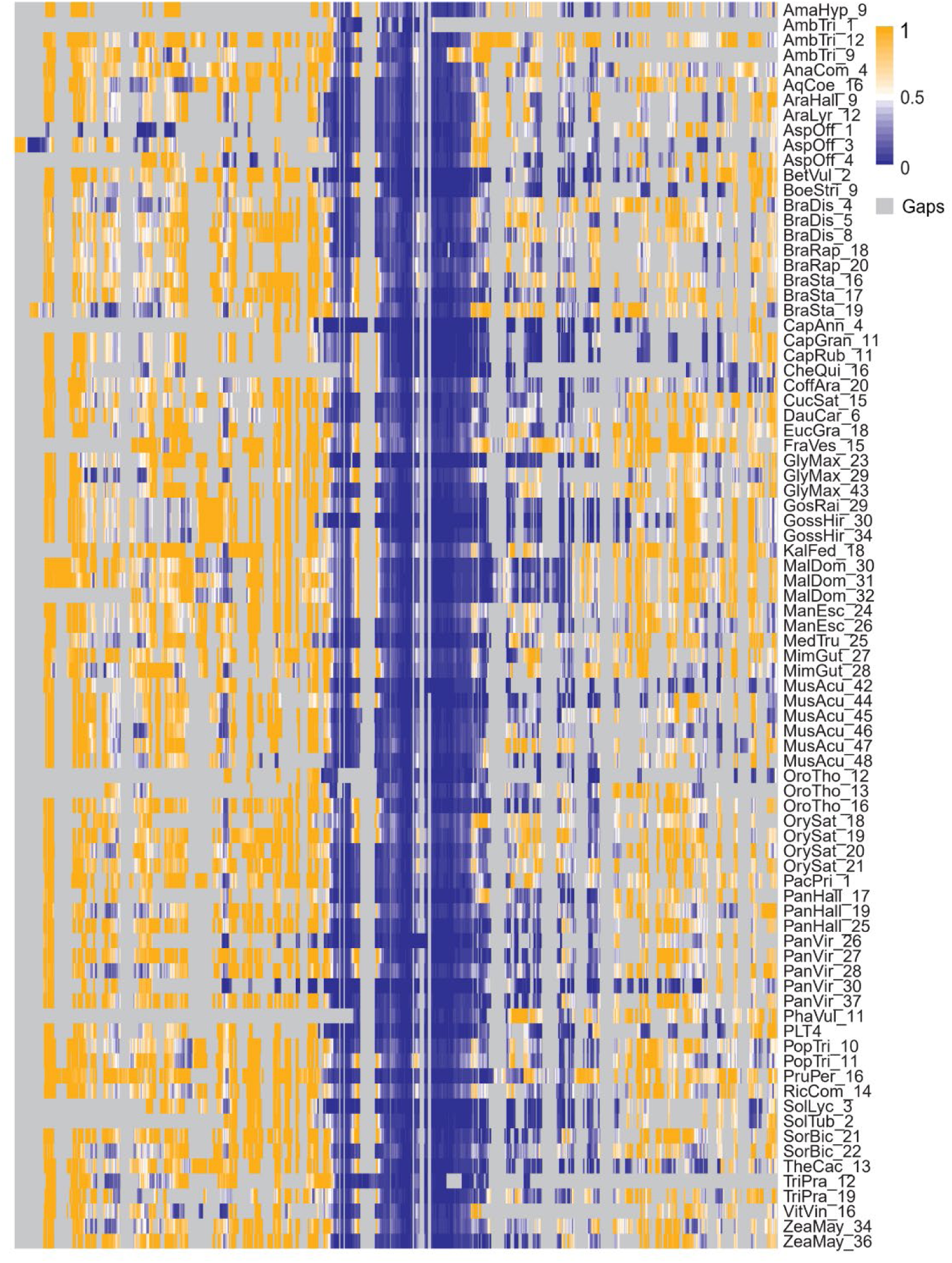
Disorder propensity of putative orthologs of *Arabidopsis thaliana* PLETHORA4. Disorder propensity was analyzed for each PLT sequence of the PLT4 (BBM) clade (Fig 2). Each amino acid in the multiple sequence alignment of the PLT4 clade was substituted by its disorder propensity value. The resulting matrix was visualized as a heatmap. Note that this analysis suggests that the AP2-linker-AP2 region, which corresponds to the DNA binding domains, is structured (blue), in line with a crystal structure obtained for a single AP2 domain of TEM1 TF (7ET4) (88). For complete species names, see S3 Table.

**S5 Fig.**
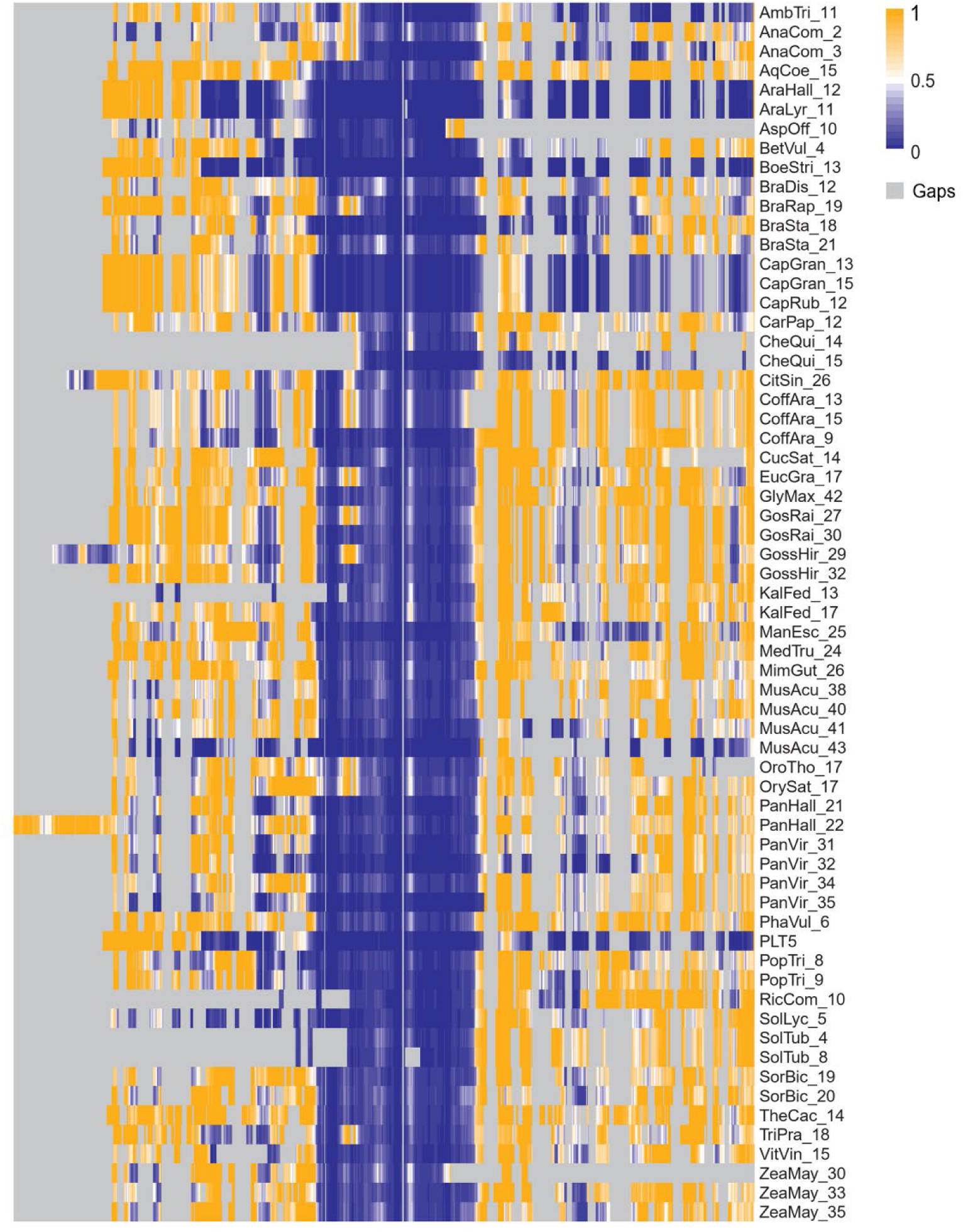
Disorder propensity of putative orthologs of *Arabidopsis thaliana* PLETHORA5. Disorder propensity was analyzed for each PLT sequence of the PLT5 clade (Fig 2). Each amino acid in the multiple sequence alignment of the proteins from the PLT5 clade was substituted by its disorder propensity value. The resulting matrix was visualized as a heatmap. Note that this analysis suggests that the AP2-linker-AP2 region, which corresponds to the DNA binding domains (will be marked in the figures), is structured (blue), in line with a crystal structure obtained for a single AP2 domain of TEM1 TF (7ET4) (88). For complete species names, see S3 Table.

**S6 Fig.**
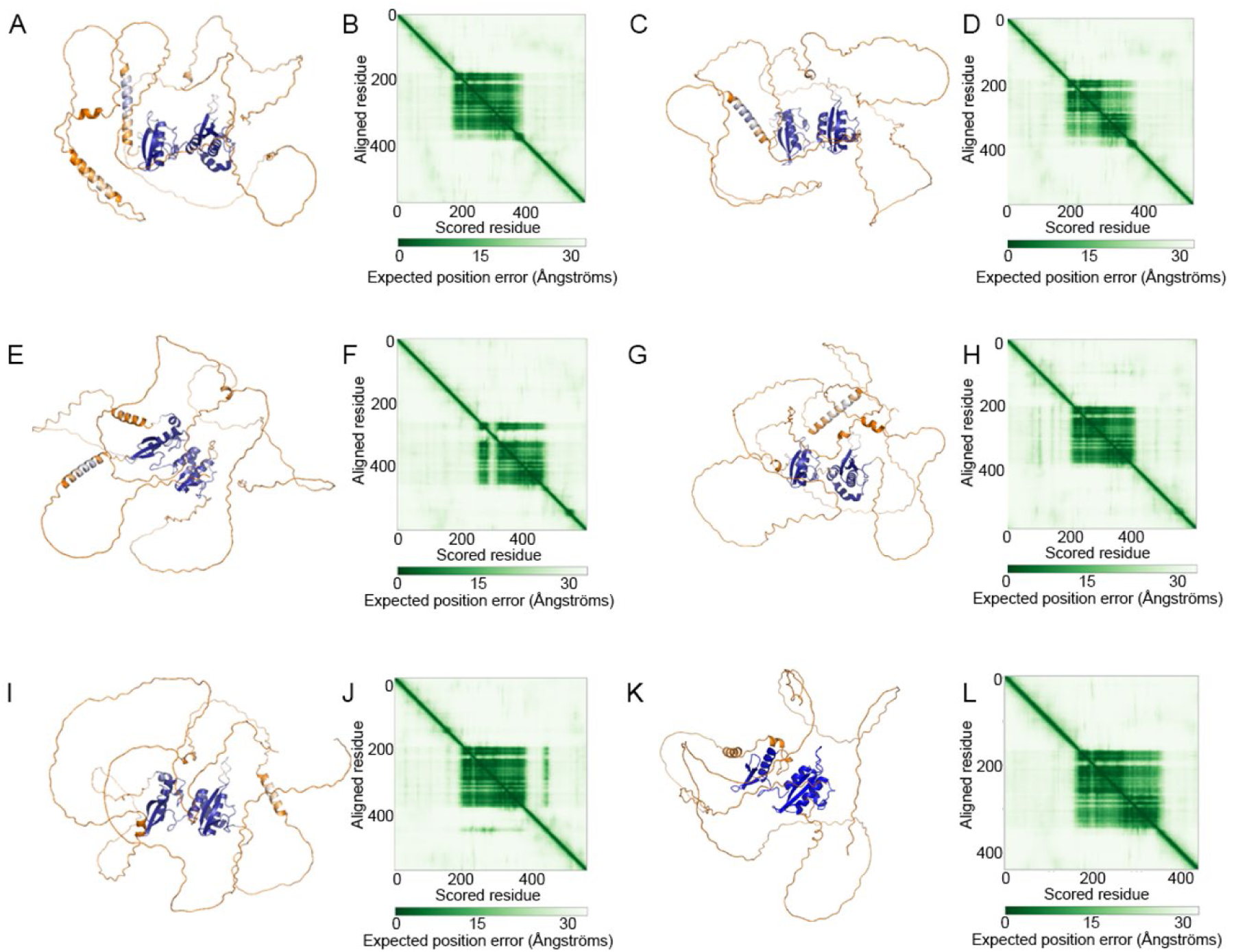
AlphaFold-predicted structures of the six PLETHORA proteins from *Arabidopsis thaliana*. A, C, E, G, I, K: Colors were assigned to the structures according to their disorder propensity values from S2-5 Figures. **B, D, F, H, J, L:** In agreement with the disorder values, AlphaFold failed to assign a putative structure to regions with higher expected position errors (yellow), supporting the notion that these might be disordered or flexible regions. **A, B:** PLT1; **C, D:** PLT2; **E, F:** PLT3; **G, H:** PLT4; **I, J:** PLT5 **L, K:** PLT7.

**S7 Fig.**
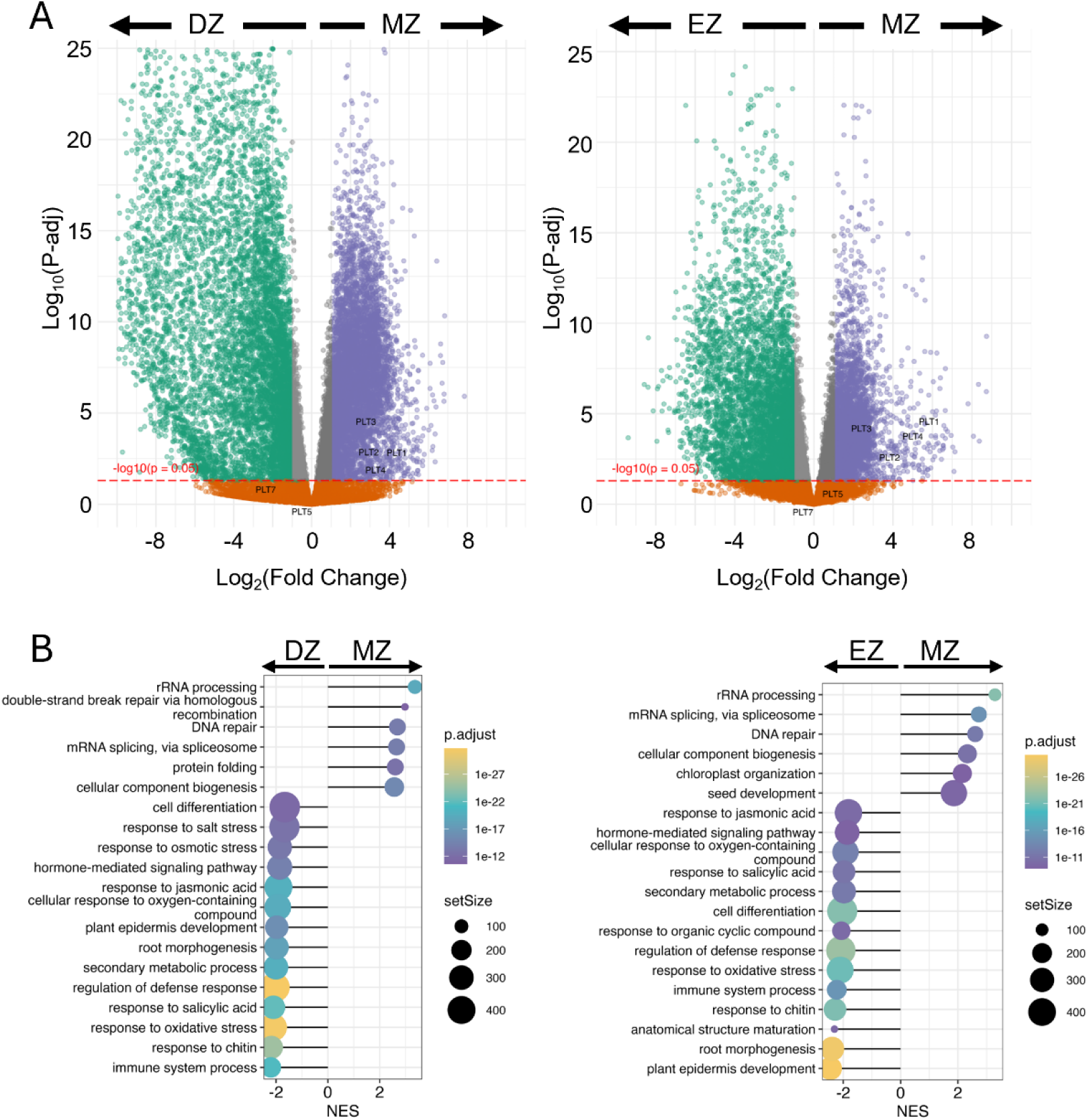
Differential gene expression and enriched biological process categories of DEGs in the *Arabidopsis thaliana* root zones. **A**. Differential gene expression analysis of the RNA-seq data from Huang and Schiefelbein, 2015 (42) performed using DESeq2, fold change ≥2. The set of upregulated genes in the meristematic zone (purple) includes most *PLETHORA* genes, as labeled in the figure. **B**. A hypergeometric test based on annotations of the differentially expressed genes in each comparison. NES = Normalized enrichment score.

**S8 Fig.**
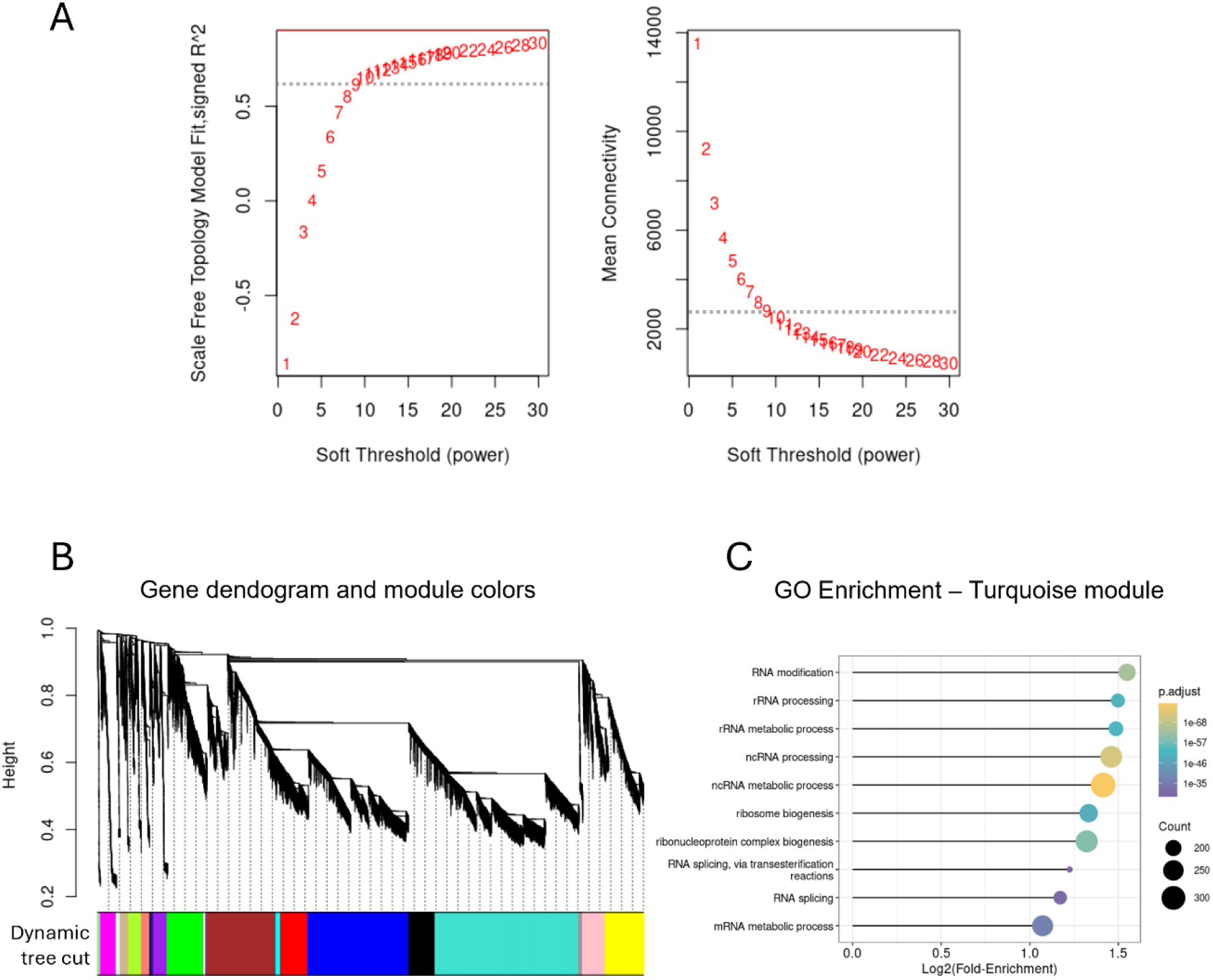
Weighted Gene Correlation Network Analysis (WGCNA) of DEGs in the *Arabidopsis thaliana* root zones. **A**. Data from *A. thaliana* meristematic, elongation, and differentiation zones of the primary root generated by Huang and Schiefelbein (42) were used for WGCNA. A soft threshold of 9 was used to ensure scale-free dependence. **B.** The turquoise module, which contains PLETHORA transcription factors, was selected for further analysis. **C**. The enrichment of biological processes represented by the nodes in the turquoise module shows an overrepresentation of rRNA and ncRNA metabolism, ribosome and ribonucleoprotein biogenesis, and RNA splicing.

**S9 Fig.**
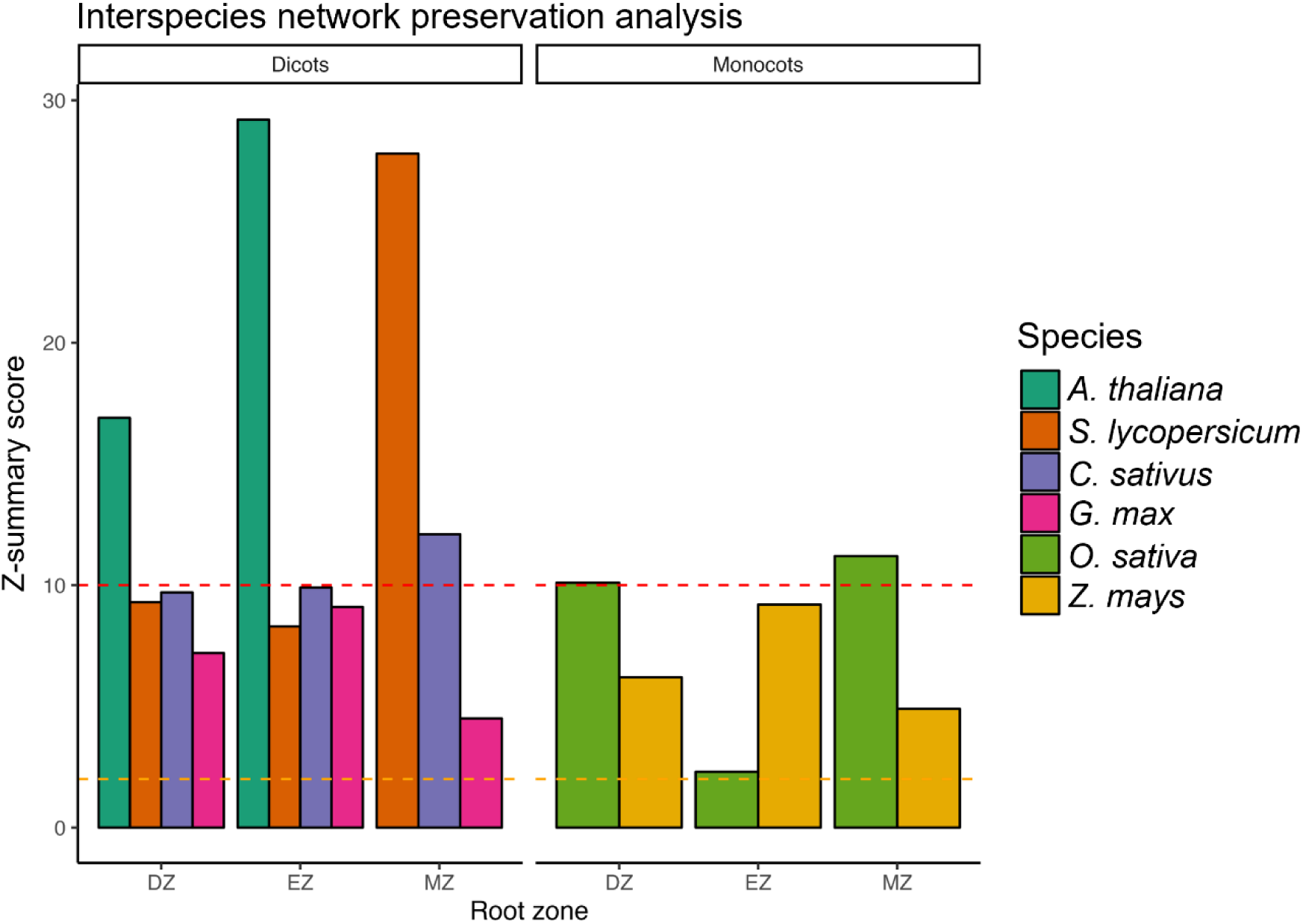
The PLETHORA-driven gene regulatory network is conserved across angiosperms. Comparisons of the connectivity and density scores of the WGCNA network for each analyzed species, presented as Z-summary scores. Values >2 suggest evidence of conservation, while values >10 suggest strong evidence of conservation according to Langfelder et al. (2011). The amino acid sequences of proteins encoded by the genes included in the *Arabidopsis thaliana* network were used as queries to search for putative orthologs in the analyzed species.

**S10 Fig.**
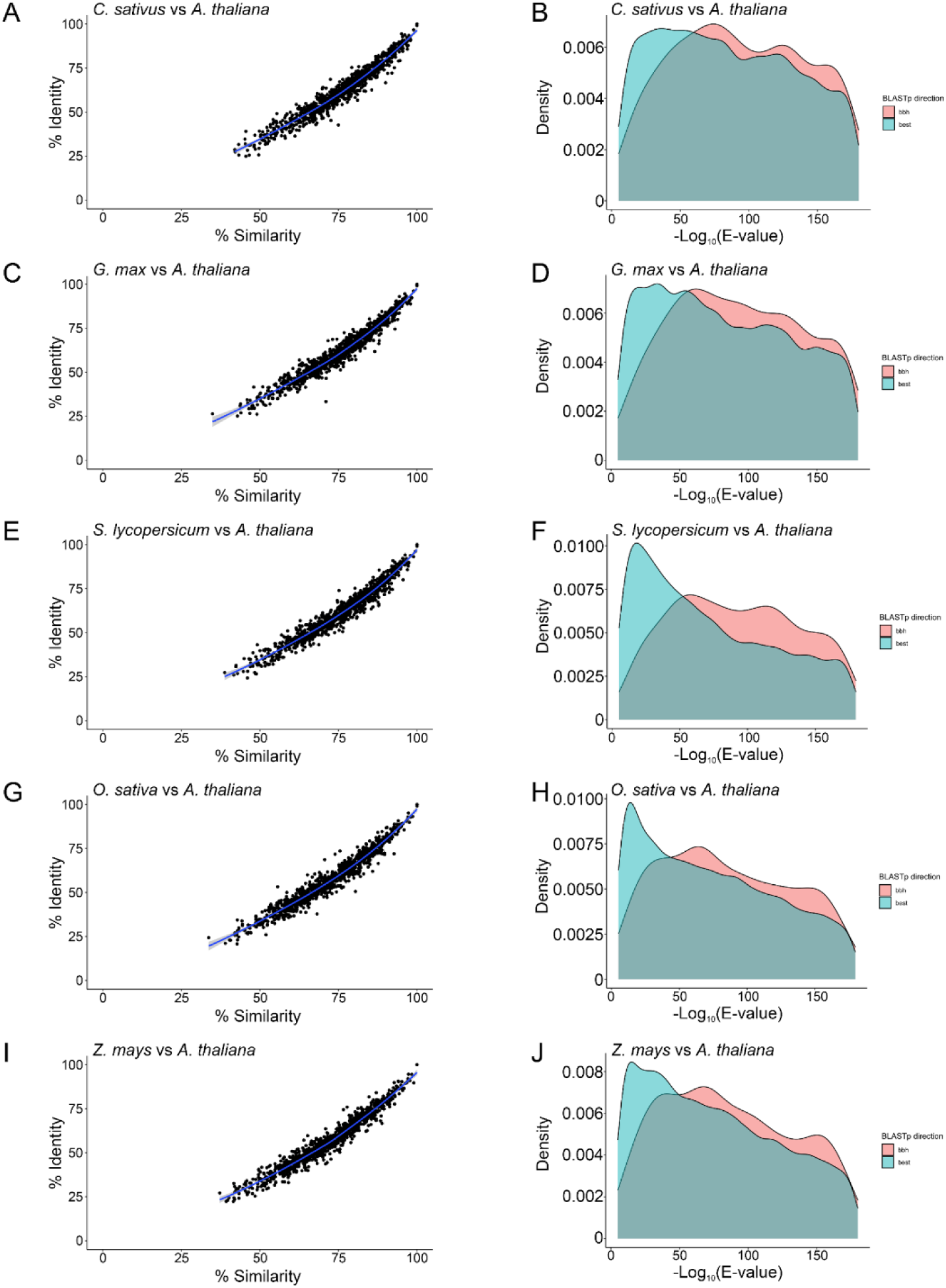
The nodes of the PLETHORA-driven gene regulatory network are highly conserved across angiosperms. The amino acid sequences of the proteins encoded by the genes in the *Arabidopsis thaliana* GRN were used as queries to search for putative orthologs in the other species. **A, C, E, G, I:** Identity and similarity scores between *A. thaliana* and putative orthologs in each species determined by pairwise BLASTp. **B, D, F, H, J:** Distribution of e-value for putative orthologs in both best bidirectional hits (bbh) and unidirectional hits (best).

**S11 Fig.**
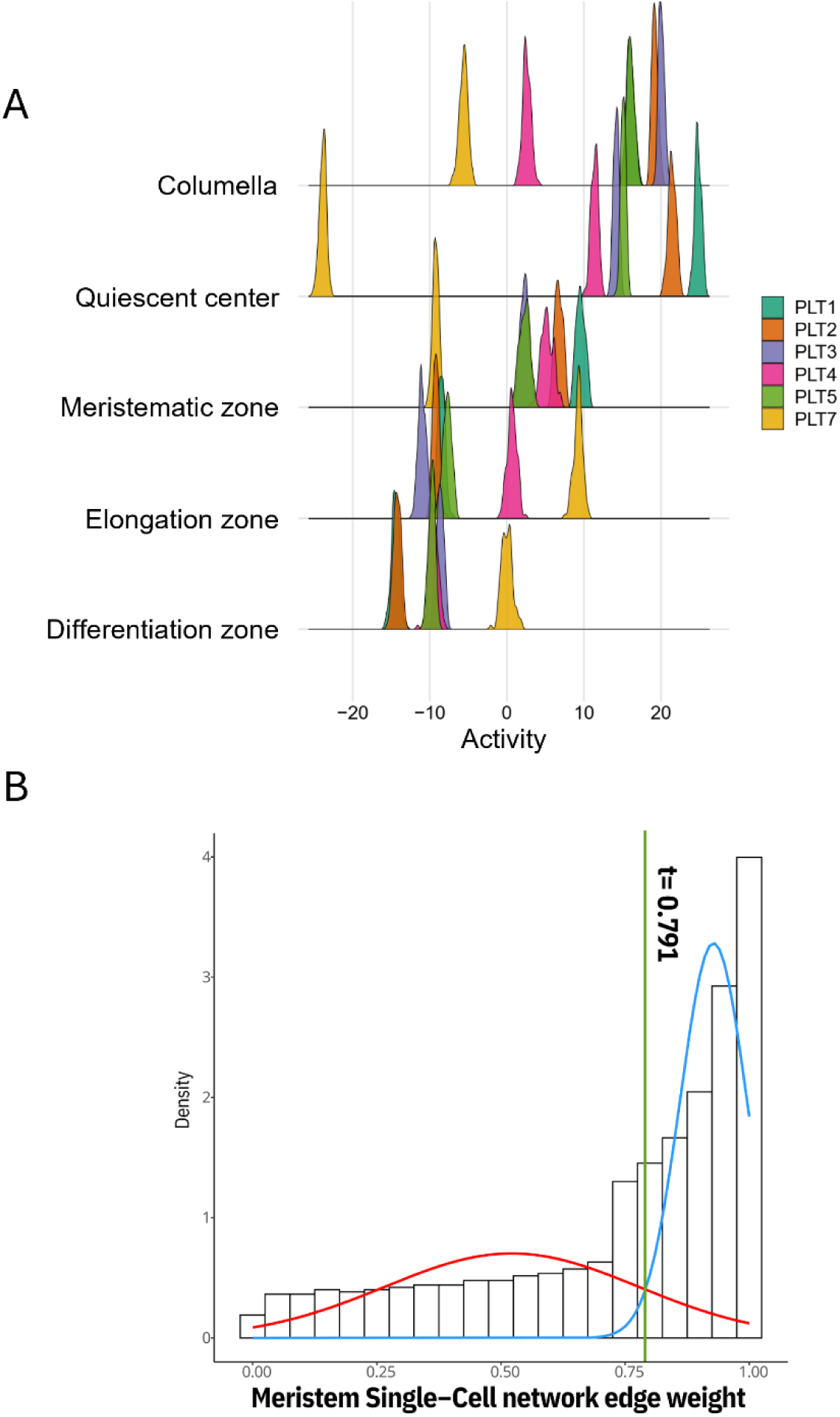
PLETHORA activity across cell types and root zones. **A.** The activity (defined as an interpolated and normalized expression profile) of each PLETHORA was calculated to allow cell-type comparisons. Note that PLT1 to PLT4, as well as PLT5, exhibit high positive activity scores in the quiescent center, with the highest score for PLT1 and PLT2. These PLTs also showed high positive activity in the root meristematic zone. **B**. Gaussian mixture model decomposition analysis to discriminate strong (blue curve) and weak (red curve) interactions in the GRN suggests a threshold of 0.79 for putative biologically relevant interactions (green line). This threshold is used in Figure 5B.

## Supplemental data

### S1 data. Identification of putative *PLETHORA* orthologs of Cactaceae

Cactaceae systematics is often challenging, and while it is somewhat robust at the subfamily level, genus and species or infraspecies relationships change frequently. This is in part because genomic resources for Cactaceae are scarce, with only two draft genomes reported: those of *Carnegiea gigantea* (Copetti et al., 2023) and *Hylocereus undatus* (now *Selenicereus undatus*) (Zheng et al., 2021). In addition, five low-coverage and highly fragmented genomes are publicly available for the species *Lophocereus schottii, Stenocereus thurberi, Pachycereus pringlei, Pereskia humboldtii* (Copetti et al., 2017), and *Cereus fernambucensis* (Amaral et al., 2021). Therefore, most molecular systematic studies of Cactaceae have been performed using plastid or mitochondrial markers, whose evolutionary rates and inheritance patterns differ from those of nuclear genes. As a consequence, assigning orthology relationships between genes is difficult and often leads to mistakenly choosing paralogs instead of orthologs across species.

To overcome this problem, we identified putative orthologs of the five PLTs previously annotated in the root apex transcriptome of *P. pringlei* (32; see S1 Table for contig IDs) in the genomes of *C. gigantea*, *L. schottii*, *S. thurberi*, *H. undatus*, and *P. pringlei* (33). All these species belong to the largest Cactaceae subfamily, Cactoideae, which includes ca. 80% of Cactaceae species; within Cactoideae, *H. undatus* is from the Hylocereeae tribe, while the four other species are members of the Pachycereeae tribe. We also compared the identity percentages of *PLT* coding sequences (CDSs).

To identify the putative *PLT* orthologs in cacti, a BLASTn search was conducted for each *P. pringlei PLT* transcript: *PLT1/2a, PLT1/2b, PLT3/7a, PLT3/7b,* and *PLT4* as queries. Hits with an e-value <1×10^-50^ were considered to be putative *PLT* orthologs. The putative 5’ and 3’ untranslated regions (UTRs) and the CDS were inferred from pairwise alignments of each scaffold with its respective *PLT* ortholog contig from the *P. pringlei* transcriptome.

The gene models of the cactus *PLT* genes are shown below: *Ppr*, *Pachycereus pringlei*; *Cgi*, *Carnegiea gigantea*; *Lsc*, *Lophocereus schottii*; *Sth*, *Stenocereus thurberi*; *Hun*, *Hylocereus undatus*. *Arabidopsis thaliana (Ath*) *PLT* genes were extracted from the Phytozome v13 database (https://phytozome-next.jgi.doe.gov/) and are shown for comparison. A 2 kilobase region upstream of the inferred putative transcription start site (TSS) is also shown as red rectangle, when available.

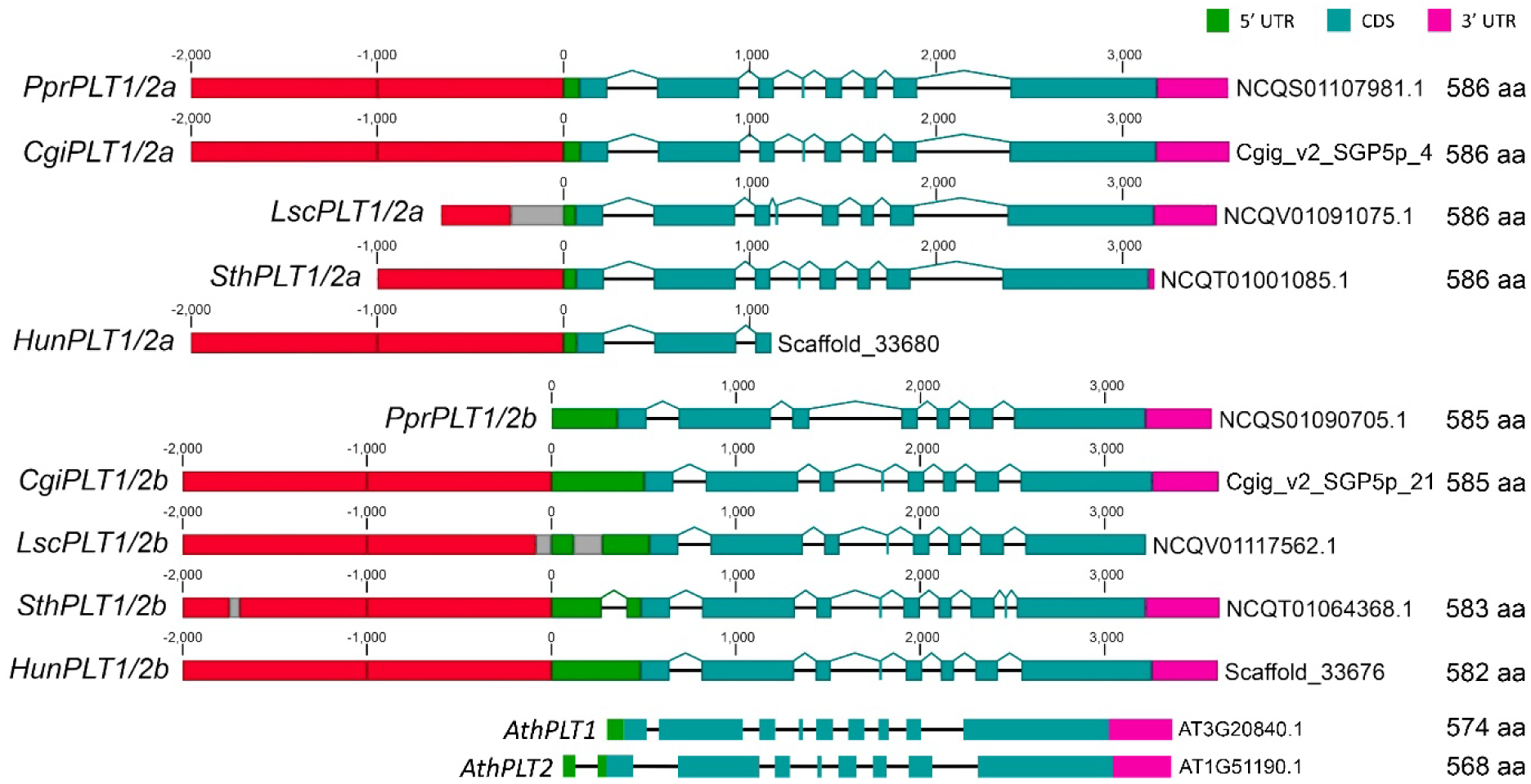

Putative Cactaceae orthologs of *PprPLT1/2a* and *PprPLT1/2b,* two genes from the PLT1-PLT2 clade, share similar exon numbers and lengths with their *Arabidopsis thaliana* counterparts. The intron lengths are also conserved in the genomic sequences of every group of Cactaceae orthologs, *PLT1/2a* and *PLT1/2b*. The scaffold identifiers and lengths of the encoded protein are shown on the right. Note that scaffolds containing some cactus genes are truncated within the coding sequence, inferred 3’-UTR, or putative promoter region before the 2 kb region. Unresolved nucleotides (Ns) are shown as gray rectangles.

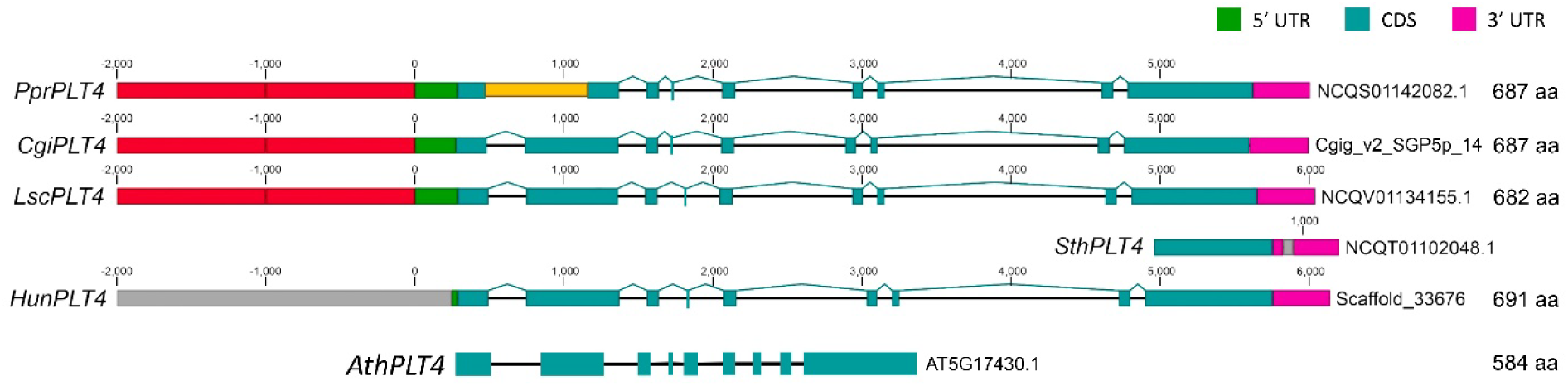

Putative cacti orthologs of *PprPLT4* share similar exon numbers and lengths with *AthPLT4*. The intron lengths are also conserved in the genomic sequences of the Cactaceae orthologs. The scaffold identifiers and lengths of the encoded protein are shown on the right. Note that in the *S. thurberi* scaffold, most of the *SthPLT4* genomic sequence is missing, while part of the *PprPLT4* genomic region failed to align with the *PprPLT4* transcript or the orthologs from all other species (yellow rectangle). The extensive region of *H. undatus* scaffold consists of the unresolved nucleotides (Ns; gray rectangle), which might be a consequence of the sequencing methodology (Illumina short reads and Hi-C).

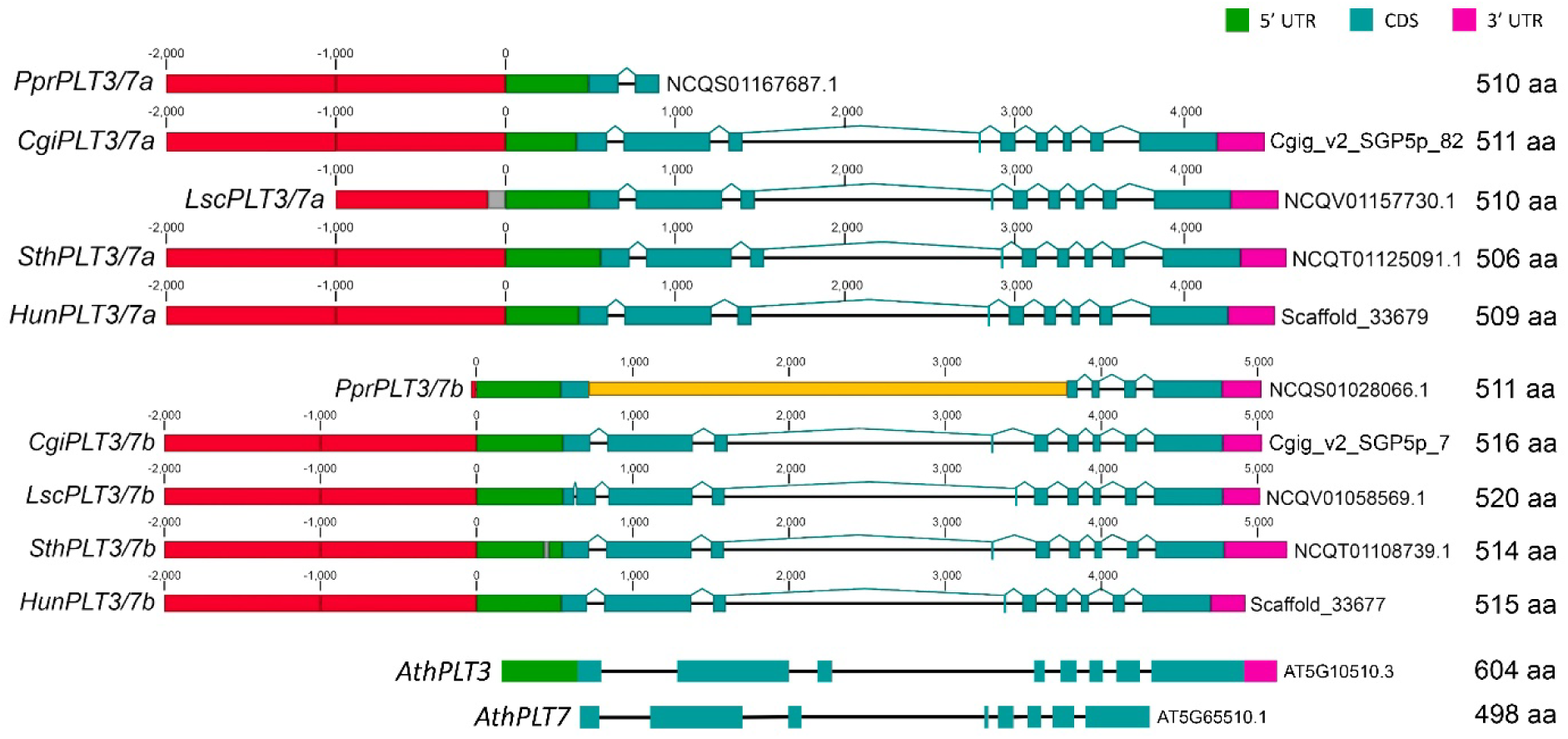

Putative Cactaceae orthologs of *PprPLT3/7a* and *PprPLT3/7b*, two genes from the PLT3 - PLT7 clade, share similar exon numbers and lengths with their counterparts. Intron lengths are also conserved in the genomic sequences of every group of Cactaceae orthologs. Scaffold identifiers and lengths of the encoded protein are shown on the right. Note that the scaffold to which *PprPLT3/7a* was mapped contains only a small region towards the 5’ of the CDS, and more than half of the *PprPLT3/7b* genomic region failed to align with the *PprPLT3/7b* transcript or the orthologs from all other species (yellow rectangle), suggesting this genomic fragment was incorrectly assembled.

Next, we analyzed the identity percentage of the CDSs of the possible Cactaceae orthologs. A BLASTn pairwise comparison revealed >94% nucleotide sequences identity; the CDS lengths of the orthologs were identical or very similar:

| *PLT1/2a* | *P. pringlei*<br>(1,761 nts) | *C. gigantea*<br>(1,761 nts) | *L. schottii*<br>(1,761 nts) |
| --- | --- | --- | --- |
| *C. gigantea*<br>(1,761 nts) | 98.6 |  |  |
| *L. schottii*<br>(1,761 nts) | 98.2 | 98.1 |  |
| *S. thurberi*<br>(1,761 nts) | 97.6 | 97.3 | 96.9 |

| **PLT1/2b** | ***P. pringlei***<br>(1,758 nts) | ***C. gigantea***<br>(1,758 nts) | ***S. thurberi***<br>(1,752 nts) |
| --- | --- | --- | --- |
| ***C. gigantea***<br>(1,758 nts) | 98.5 |  |  |
| ***S. thurberi***<br>(1,752 nts) | 95.0 | 94.4 |  |
| ***H. undatus***<br>(1,749 nts) | 95.5 | 94.8 | 95.7 |

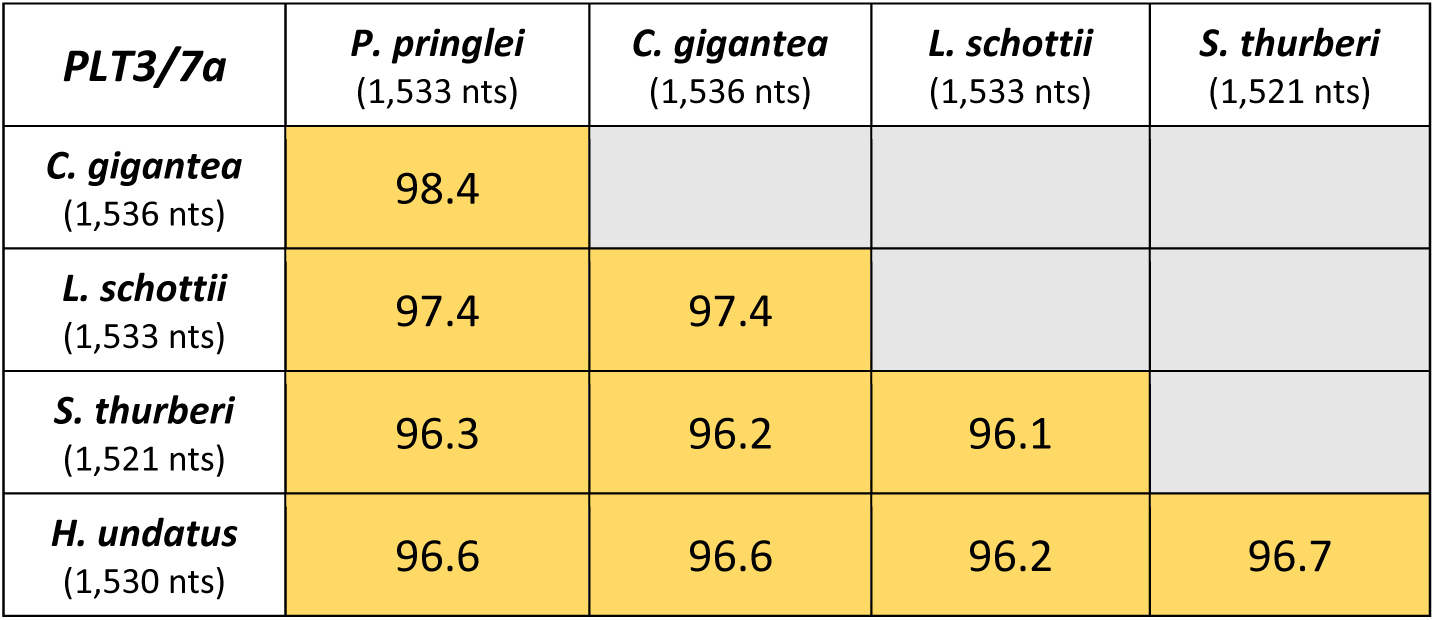

| **PLT3/7b** | ***P. pringlei***<br>(1,536 nts) | ***C. gigantea***<br>(1,551 nts) | ***L. schottii***<br>(1,563 nts) | ***S. thurberi***<br>(1,545 nts) |
| --- | --- | --- | --- | --- |
| ***C. gigantea***<br>(1,551 nts) | 97.4 |  |  |  |
| ***L. schottii***<br>(1,563 nts) | 96.3 | 95.4 |  |  |
| ***S. thurberi***<br>(1,545 nts) | 97.1 | 97.1 | 95.7 |  |
| ***H. undatus***<br>(1,548 nts) | 95.6 | 96.0 | 94.3 | 96.5 |

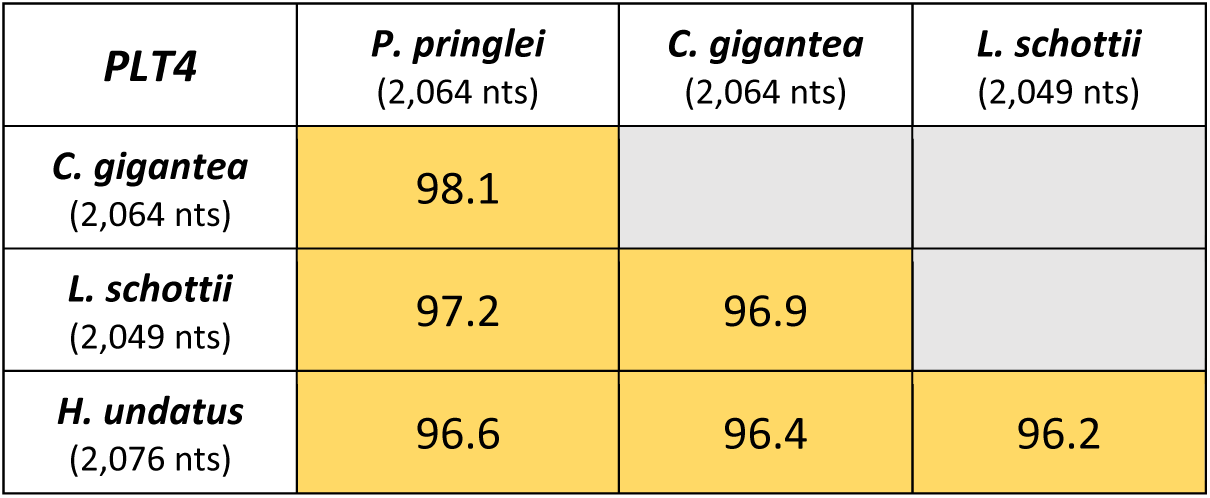

| **PLT4** | ***P. pringlei***<br>(2,064 nts) | ***C. gigantea***<br>(2,064 nts) | ***L. schottii***<br>(2,049 nts) |
| --- | --- | --- | --- |
| ***C. gigantea***<br>(2,064 nts) | 98.1 |  |  |
| ***L. schottii***<br>(2,049 nts) | 97.2 | 96.9 |  |
| ***H. undatus***<br>(2,076 nts) | 96.6 | 96.4 | 96.2 |

By contrast, the CDS and intron lengths of the PLT paralogs were different, and their identity percentages were much smaller, as shown here for the CDS and intron lengths *Carnegiea gigantea* paralogs (CDS length is shown on the right):

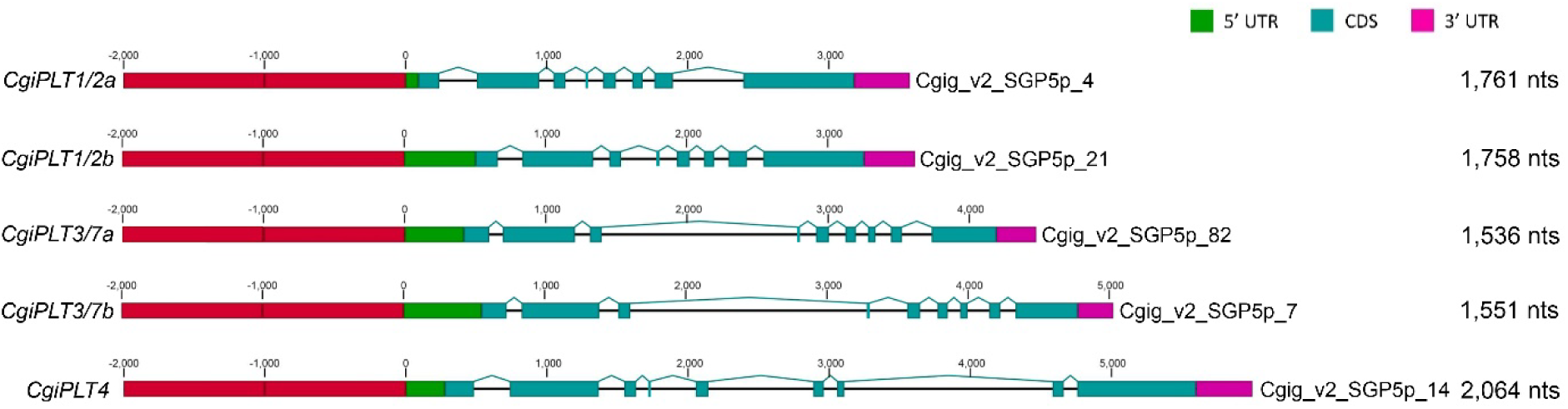

and CDS identity percentages of *Pachycereus pringlei* PLT paralogs (*de novo* assembled transcripts (32).

|  | *PprPLT1/2a*<br>(1,761 nts) | *PprPLT1/2b*<br>(1,758 nts) | *PprPLT3/7a*<br>(1,533 nts) | *PprPLT3/7b*<br>(1,536 nts) |
| --- | --- | --- | --- | --- |
| *PprPLT1/2b*<br>(1,758 nts) | 68.8 |  |  |  |
| *PprPLT3/7a*<br>(1,533 nts) | 50.3 | 50.0 |  |  |
| *PprPLT3/7b*<br>(1,536 nts) | 50.5 | 50.8 | 73.5 |  |
| *PprPLT4*<br>(2,064 nts) | 51.5 | 53.4 | 45.1 | 44.8 |

Therefore, the CDSs of all putative *PLT* orthologs of five Cactaceae species have similar lengths and share more than 94% identity, while the lengths of the CDSs of the *PLT* paralogs are variable, with identity below 75%.

In conclusion, in the absence of well-assembled genomes (which hindered our implementation of synteny or microsynteny analysis), we showed that gene models could be reconstructed from the available data, including inferring UTRs, intronic regions, and exonic regions, to better assess the orthology relationships of non-model species. In the case of PLTs, which are a particularly difficult to analyze given the duplication of PLT family members, this approach overcomes the need to reconstruct the phylogeny, which could otherwise introduce bias and might mistakenly group sequences due to the inclusion of incomplete protein sequences.

## **Supplemental tables** (uploaded file “*Sup_Tables.xlsx*”)

**S1 Table. List of AP2 protein sequences from Embryophyta genomes.** All sequences reported in this table contain two AP2 domains separated by an interdomain sequence.

**S2 Table. Evolutionary model selection for phylogeny reconstruction**. The model selected for multiple sequence alignment of both whole protein sequences and the AP2- interdomain-AP2 region was the Jones-Taylor-Thornton (JTT) model with four gamma rate categories (+G4). For the AP2-interdomain-AP2 region, the selected model also included estimated amino acid frequencies (+F). This model is thought to be the most robust based on the Bayesian information criterion (BIC), Akaike information criterion (AIC), and corrected AIC (AICc).

**S3 Table. List of sequence identifiers of putative PLT proteins.** The database IDs are listed in S1 Table.

**S4 Table. Differentially expressed genes in paired comparisons between the meristematic (MZ), elongation (EZ), and differentiation (DZ) root zones of selected species.**

**S5 Table. Interactions included in the PLT-regulated network**.

**S6 Table. Properties of nodes in the PLT-regulated network**. Columns ending in _rep1, 2, or 3 list the RPKMs of the genes in each biological replicate. Raw data were obtained from Huang and Schiefelbein, 2015 (42).

## Author Contributions

Conceptualization and study design: GRA, SS; Formal analysis: JRH, KAGA, RUAH, JPVN, GRA, SS; Funding acquisition: SS; Supervision, GRA, SS; Writing – original draft: GRA, SS; Writing – review and editing: JRH, KAGA, JPVN, GRA, SS.

## Acknowledgements

We thank A. Hetheringon and Yanbin Yin for granting access to the *Isoetes echinospora* transcriptome and *Hylocereus/Selenicereus undatus* genome, respectively; S. Mapsucialy-Mendivil, J.M. Hurtado Ramírez, S. Ainsworth, for technical help; J.F. Martínez-Rodríguez and F. Molina-Freaner for *Pachycereus pringlei* and *Carnegiea gigantea* seed donation.

GRA acknowledges postdoctoral fellowship from Consejo Nacional de Ciencia y Tecnología (CONACyT); RUAH and JPVN thanks the support of CONACyT through a MSc Scholarships (CVU 997135 and 1074303, respectively).

